# DNA Structure Design Is Improved Using an Artificially Expanded Alphabet of Base Pairs Including Loop and Mismatch Thermodynamic Parameters

**DOI:** 10.1101/2023.06.06.543917

**Authors:** Tuan M. Pham, Terrel Miffin, Hongying Sun, Kenneth K. Sharp, Xiaoyu Wang, Mingyi Zhu, Shuichi Hoshika, Raymond J. Peterson, Steven A. Benner, Jason D. Kahn, David H. Mathews

## Abstract

We show that *in silico* design of DNA secondary structures is improved by extending the base pairing alphabet beyond A-T and G-C to include the pair between 2-amino-8-(1’-β-D-2’-deoxyribofuranosyl)-imidazo-[1,2-*a*]-1,3,5-triazin-(8*H*)-4-one and 6-amino-3-(1’-β-D-2’-deoxyribofuranosyl)-5-nitro-(1*H*)-pyridin-2-one, simply P and Z. To obtain the thermodynamic parameters needed to include P-Z pairs in the designs, we performed 47 optical melting experiments and combined the results with previous work to fit a new set of free energy and enthalpy nearest neighbor folding parameters for P-Z pairs and G-Z wobble pairs. We find that G-Z pairs have stability comparable to A-T pairs and therefore should be considered quantitatively by structure prediction and design algorithms. Additionally, we extrapolated the set of loop, terminal mismatch, and dangling end parameters to include P and Z nucleotides. These parameters were incorporated into the RNAstructure software package for secondary structure prediction and analysis. Using the RNAstructure Design program, we solved 99 of the 100 design problems posed by Eterna using the ACGT alphabet or supplementing with P-Z pairs. Extending the alphabet reduced the propensity of sequences to fold into off-target structures, as evaluated by the normalized ensemble defect (NED). The NED values were improved relative to those from the Eterna example solutions in 91 of 99 cases where Eterna-player solutions were provided. P-Z-containing designs had average NED values of 0.040, significantly below the 0.074 of standard-DNA-only designs, and inclusion of the P-Z pairs decreased the time needed to converge on a design. This work provides a sample pipeline for inclusion of any expanded alphabet nucleotides into prediction and design workflows.

## INTRODUCTION

Natural and designed nucleic acids serve a number of roles *in vitro* and in cells. In nature, DNA is largely an information carrier, but RNA is an information carrier, a catalyst,^1^ an agent for recognition of complementary sequences, and an aptamer for metabolite binding.^2–4^ New binding and catalytic roles can be evolved *in vitro* for RNA and DNA using systematic evolution of ligands by exponential enrichment (SELEX).^5^ RNA and DNA are also convenient scaffolds for molecular arrays^6, 7^ and molecular machines.^8^ Almost all of these roles for nucleic acids involve folding into secondary and tertiary structures beyond simple duplexes.

Nucleic acids can be designed *in silico* to fold into specified secondary structures, and a reliable method for solving this so-called inverse folding problem would reduce the need for selection or trial and error experiments.^9^ Traditionally, the design process requires a search algorithm and an objective function to evaluate candidate designs. An efficient search algorithm is necessary because] an exponentially increasing number of sequences] can fold to any desired target structure as a function of sequence length.^10^ Generally, sequences are optimized locally and subsequently assembled into larger structures. The objective function assesses the design quality by comparison to the target structure.

One approach is to guarantee that the sequence folds with minimum (most negative) free energy change from the random coil to give the target structure.^11^ This ensures that the most populated structure at equilibrium will be the target structure, although it does not guarantee that the probability of the desired structure will be high in the Boltzmann ensemble.

Another approach is to minimize the normalized ensemble defect (NED), the average probability that a nucleotide will be the wrong conformation in the ensemble.^12^ This improves upon assessing solely the minimum free energy structure in that optimizing the NED also ensures that the helix and loop components of the desired structure will also occur with high probability.^13^ It can conveniently be calculated from a partition function expression^14^ for base pairing probabilities:

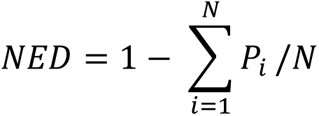

where *N* is the sequence length and *P_i_* is the probability that nucleotide *i* is in the expected structure. The probability *P_i_* is computed over the Boltzmann ensemble of all possible secondary structures.^14^ For paired nucleotides, *P_i_* is the probability that nucleotide *i* is paired to its exact intended pairing partner. For unpaired nucleotides, it is the probability the nucleotide is unpaired.

The normalized ensemble defect quantifies the extent to which the desired components of the structure dominate the Boltzmann ensemble. It can be difficult to achieve a low ensemble defect because many competing structures are often possible for a sequence. The desired structure and its close analogs must have a substantially lower folding free energy change than all of the competing structures in order to fold with high probability.

In this work, we design structures with an extended DNA alphabet including the P (Imidazo-[1,2-*a*]-1,3,5-triazin-(8*H*)-4-one) and Z (6-Amino-5-nitro-(1*H*)-pyridin-2-one) bases of the second-generation AEGIS (Artificially Expanded Genetic Information System). P and Z form a three hydrogen-bonded base pair (Fig. 1A)^15, 16^ that is largely orthogonal to the standard Watson-Crick-Franklin base pairs. Hence, P and Z can be used to add base pairs that are less likely to find alternative pairing partners than natural DNA nucleotides.

**Figure 1.**
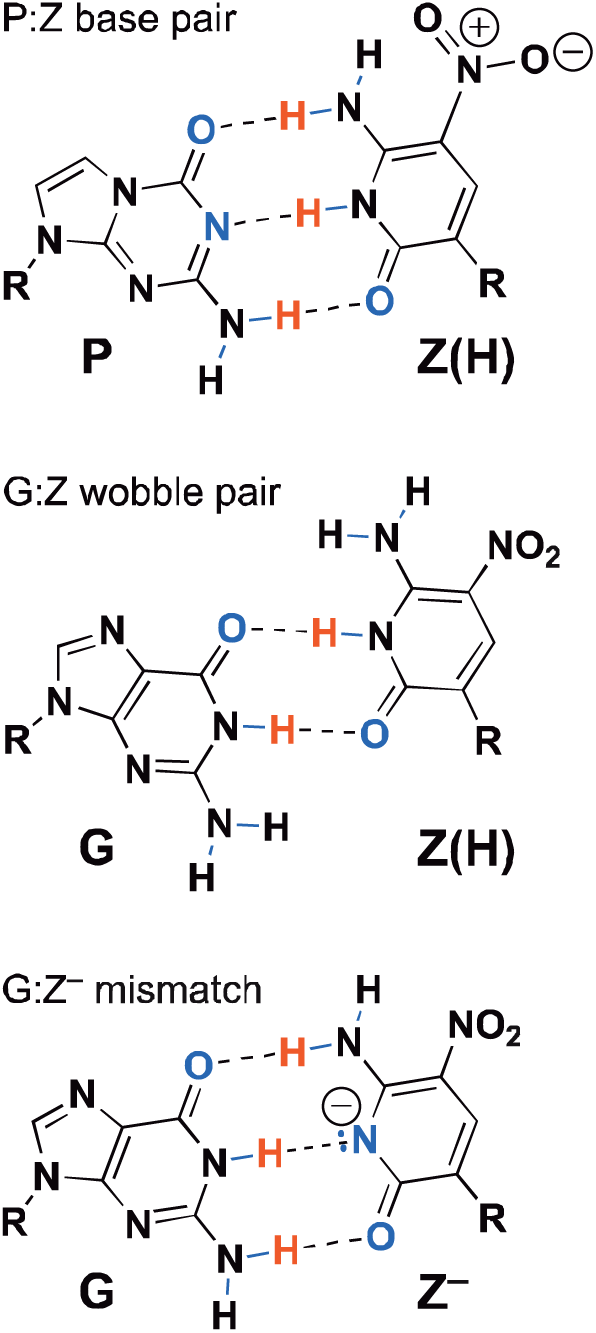
(Top) The P-Z base pair has three hydrogen bonds. (Middle) The proposed G-Z “wobble” base pair with two hydrogen bonds. (Bottom) The deprotonated G-Z^−^ pair is dominant at elevated pH.^17^

To use P and Z in inverse folding designs, the thermodynamics of folding from the random coil need to be known in order to evaluate the design quality. Prior work established that P and Z prefer pairing with each other above all single mismatches, except that G-Z pairs are roughly as stable as A-T pairs (Fig. 1).^16, 17^ A subsequent study fit the helical nearest neighbor thermodynamic parameters for P-Z as part of the eight base “Hachimoji” system.^15^ In this work, we determined nearest neighbor parameters for a full P-Z extended alphabet including fitting stacks with G-Z wobbles, better defining the P-Z stack parameter values, and extrapolating loop parameters to include P and Z bases. The RNAstructure software package was previously extended to accommodate folding alphabets of any size, and therefore it is able to handle designs using the six base alphabet.^18^

We demonstrate that *in silico* designs are improved using the additional P-Z base pair. We chose to use the Eterna100 benchmark structures^19^ and the design program from RNAstructure,^20^ which implements a version of the NUPACK algorithm for NED minimization.^12^ 94 of 100 benchmark structures could be designed by RNAstructure using both standard DNA and the extended P-Z base pairs. The sequences with P-Z outperformed the sequences with standard DNA, as quantitated by showing a significantly lower NED. Both types of designs had significantly lower NED than the standard solutions provided by Eterna.

## RESULTS

For optimal designs it is essential to have as complete a set of nearest neighbor thermodynamic parameters as possible to estimate folding stability from the random coil. Previous work showed that G-Z pairs stabilize helix formation^16^ and secondary structure designs have loop regions that connect helices. Therefore, we extended the existing data on P and Z thermodynamics and fit additional nearest neighbor parameters.

### Stacking Nearest Neighbor Parameters for P-Z and G-Z base pairs

To fit nearest neighbor stacking parameters for P-Z pairs, we used the existing dataset of optical melting data for duplexes with P-Z pairs^15, 16^ and an additional 13 optical melting experiments from this work. We refined the data analysis procedure for optical melting experiments to simultaneously fit absorbance vs temperature curves at multiple concentrations in order to minimize the error with respect to all available experimental data (see Materials and Methods for details). Supplemental Table S1 provides the optical melting stabilities determined for the 13 all-helix duplexes studied in this work.

An important consideration is whether terminal P-Z pairs need an end correction term like A-T pairs or RNA A-U pairs.^21, 22^ The end correction term was originally motivated by hydrogen bond counting. Subsequent studies, however, found that G-U and m^6^A-U pairs at helix ends did not need corrections,^18, 23^ which indicates that hydrogen bond counting does not determine the need for an end correction. It was therefore an open question as to whether an end correction would be necessary for P-Z pairs at helix ends. To address this, we used linear regression to fit the P-Z stacking parameters both with and without an end correction. Supplemental Table S2 shows the results of these regressions, which demonstrated that the P-Z end correction was small in magnitude compared to the standard error of the regression (0.33 ± 0.21 kcal/mol).

We examined the residuals for the fit when the P-Z end penalty was included and observed a large residual of 2.22 kcal/mol for the duplex (ZGCATGCP)_2_. This single experimental value was driving much of the magnitude of the P-Z penalty, and the residual for (ZGCATGCP)_2_ was the largest across all duplexes used in the fit. Therefore, we removed this sequence from the fit as an outlier and repeated the regression. In this fit, absent the outlier, the terminal P-Z parameter is smaller in magnitude than the standard error of the regression (0.03 ± 0.19). Given this result, we do not include a terminal P-Z penalty in the fit of the stacking nearest neighbor parameters. The additional experimental data available in this study informed this decision, which differs from previous work that did include the terminal P-Z penalty.^15^

The P-Z stack nearest neighbor parameters for free energy change at 37 °C are provided in Table 1, where they are compared to G-C nearest neighbor parameters. On average, a P-Z substitution for a G-C pair is stabilizing by −0.2 ± 0.4 kcal/mol. This varies across the stacks; there are two cases where P-Z substitutions of G-C pairs were unfavorable. Therefore, the full nearest neighbor treatment of P-Z pairs is required to fit the set of oligonucleotide thermodynamic data; a simple increment for P-Z substitution of G-C pairs would not suffice.

**Table 1.** P-Z stack nearest neighbor parameters, as compared to analogous G-C stacks (replacing P-Z with G-C). On average, the P-Z stacks are more stable by −0.2 ± 0.4 kcal/mol per P-Z pair. The free energy increment for terminal P-Z pairs is set to 0 kcal/mole, as discussed in the text.

| Parameter | $\Delta G^\circ_{37}$ (kcal/mol) | Parameter | $\Delta G^\circ_{37}$ (kcal/mol) | $\Delta\Delta G^\circ_{37}$ (kcal/mol)<br>per P-Z<br>substitution |
| --- | --- | --- | --- | --- |
| AZ<br>TP | $-1.44 \pm 0.17$ | AC<br>TG | -1.4 | 0.0 |
| TP<br>AZ | $-1.45 \pm 0.15$ | TG<br>AC | -1.5 | 0.1 |
| TZ<br>AP | $-1.49 \pm 0.15$ | TC<br>AG | -1.3 | -0.2 |
| PZ<br>ZP | $-1.57 \pm 0.33$ | GC<br>CG | -2.2 | +0.3 |
| GZ<br>CP | $-1.74 \pm 0.16$ | GC<br>CG | -2.2 | +0.5 |
| AP<br>TZ | $-1.85 \pm 0.16$ | AG<br>TC | -1.3 | -0.6 |
| CZ<br>GP | $-2.07 \pm 0.13$ | CC<br>GG | -1.8 | -0.3 |
| GP<br>CZ | $-2.28 \pm 0.16$ | GG<br>CC | -1.8 | -0.5 |
| PP<br>ZZ | $-2.35 \pm 0.13$ | GG<br>CC | -1.8 | -0.3 |
| CP<br>GZ | $-2.77 \pm 0.16$ | CG<br>GC | -2.2 | -0.6 |
| ZP<br>PZ | $-3.35 \pm 0.32$ | CG<br>GC | -2.2 | -1.2 |

Figure 2 is a plot of predicted ΔG°_37_ versus experimental ΔG°_37_ for all of the melting experiments, and the full table of residuals is provided in Supplemental Table S3. The sequence (GTPPZZAC)_2_ has the largest residual, with a predicted overstabilization by 1.41 kcal/mol. This sequence has four P-Z pairs in a row, and the residual might indicate a non-nearest-neighbor effect. (GAZZPPTC)_2_, however, also has four P-Z pairs in a row, but it has more accurately predicted stability, i.e., the residual is 0.77 kcal/mol understabilized by the prediction. Additional experiments on related sequences could be used in the future to better understand why (GTPPZZAC)2 has the largest residual and whether this reflects possible non-nearest neighbor effects of multiple adjacent P-Z pairs.

**Figure 2.**
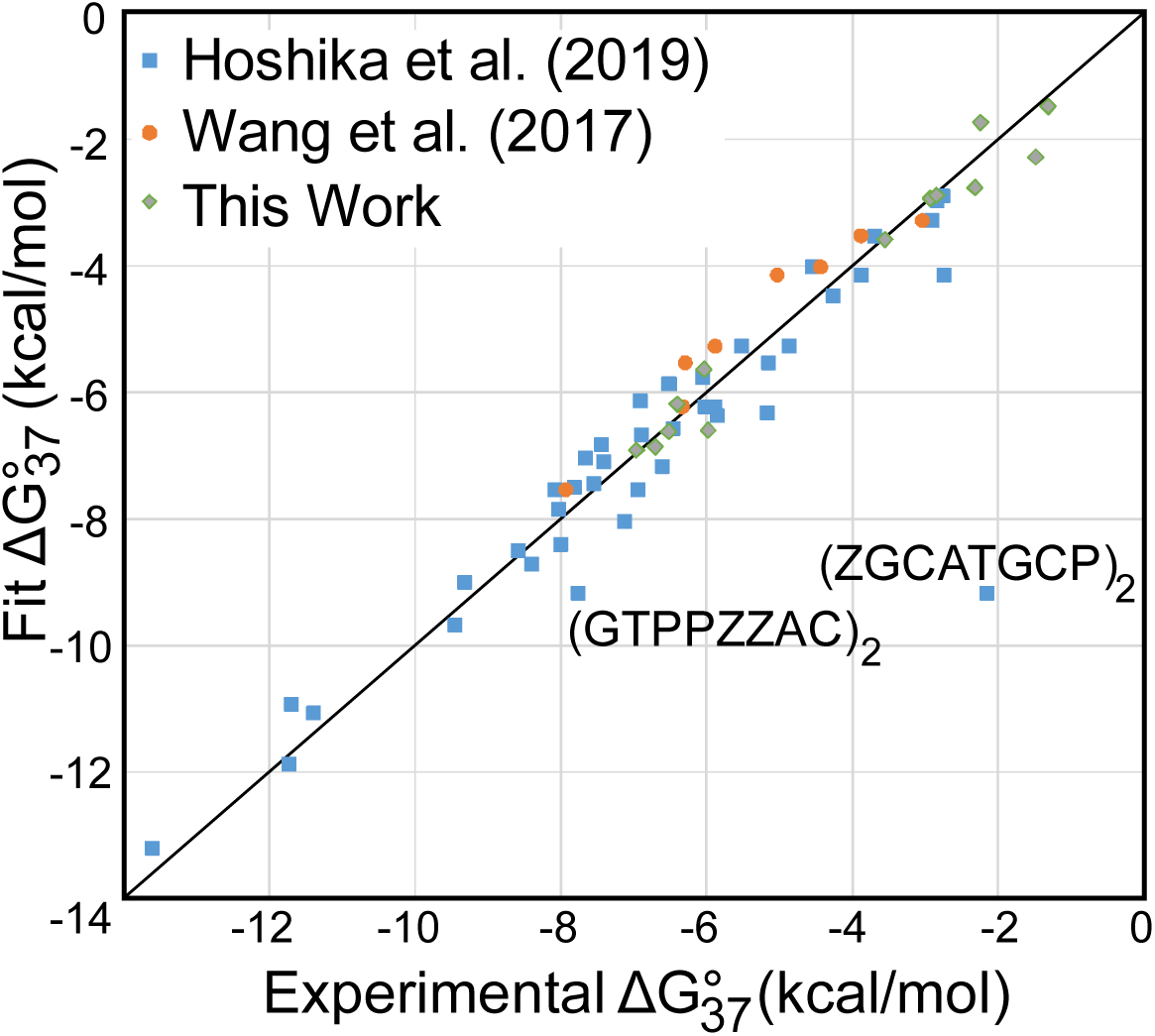
Predicted ΔG°_37_ as a function of experimental ΔG°_37_ for the P-Z stacks. The P-Z stack component is the measured folding free energy change minus the contributions of Watson-Crick-Franklin stacks, intermolecular initiation, and the symmetry correction (applied to self-complementary duplexes only). The *y* = *x* line is shown for reference. The duplex (ZGCATGCP)_2_ was removed from the fit as an outlier. The duplex (GTPPZZAC)_2_ has the largest residual for a value used in the fit, at 1.41 kcal/mol. The data were derived from Hoshika et al.,^15^ Wang et al.,^16^ and additional duplexes reported here (Table S1).

Our prior work studying the stability of P-Z pairs and mismatches with P or Z demonstrated that G-Z pairs are more stabilizing that A-T pairs, suggesting that they should be included as a wobble pair and included in secondary structure prediction, like G-U wobbles in RNA.^23^ The G-Z pair could also be a deprotonated three hydrogen-bond pair, as observed at higher pH; the thermodynamics alone at pH 7 cannot distinguish structures.^17, 24^ To fit the 15 nearest neighbor parameters for G-Z stacking on A-T, G-C, P-Z, or G-Z pairs, we performed an additional 11 melts of duplexes containing G-Z pairs and combined this with 4 prior experiments.^16^ Supplementary Table S4 provides the results of these optical melting experiments.

Table 2 provides the G-Z stack nearest neighbor parameters for free energy change at 37 °C, which were fit by linear regression (see Materials and Methods) using fixed values of stacks with P-Z pairs (Table 1). Table 2 also compares the G-Z pair stacks to A-T pair stacks. The number of sequences studied by optical melting is equal to the number of G-Z stack parameters, and therefore the error estimates, which are the standard errors of the regression, are underestimates. Supplementary Table S5 provides the residuals from the fit of the parameters.

**Table 2.** G-Z stack nearest neighbor parameters, as compared to analogous A-T stacks (replacing G-Z with A-T). The error estimates are the standard errors of the regression, but these are underestimates.

| Parameter | $\Delta G^{\circ}_{37}$<br>(kcal/mol) | Parameter | $\Delta G^{\circ}_{37}$ (kcal/mol) | $\Delta\Delta G^{\circ}_{37}$<br>(kcal/mol) per P-Z substitution |
| --- | --- | --- | --- | --- |
| GZ<br>ZG | $1.70 \pm 0.48$ | AT<br>TA | -0.9 | 2.6 |
| GA<br>ZT | $0.33 \pm 0.27$ | AA<br>TT | -1.0 | 0.9 |
| GG<br>ZZ | $0.26 \pm 0.14$ | AA<br>TT | -1.0 | 1.3 |
| CZ<br>GG | $-0.17 \pm 0.26$ | CT<br>GA | -1.3 | 0.4 |
| GP<br>ZZ | $-0.50 \pm 0.28$ | AP<br>TZ | -1.9 | 1.1 |
| AG<br>TZ | $-0.58 \pm 0.14$ | AA<br>TT | -1.0 | 0.4 |
| GG<br>CZ | $-0.75 \pm 0.20$ | GA<br>CT | -1.3 | 1.3 |
| AZ<br>TG | $-1.16 \pm 0.10$ | AT<br>TA | -0.9 | -0.7 |
| TG<br>AZ | $-1.40 \pm 0.24$ | TA<br>AT | -0.6 | 0.6 |
| ZG<br>GZ | $-1.67 \pm 0.41$ | TA<br>AT | -0.6 | -0.1 |
| CG<br>GZ | $-1.79 \pm 0.26$ | CA<br>GT | -1.5 | 0.5 |
| GZ<br>ZP | $-1.81 \pm 0.28$ | AZ<br>TP | -1.5 | -0.3 |
| ZP<br>GZ | $-1.85 \pm 0.20$ | TP<br>AZ | -1.5 | -1.1 |
| GZ<br>CG | $-2.07 \pm 0.17$ | GT<br>CA | -1.4 | -0.7 |
| PG<br>ZZ | $-2.47 \pm 0.24$ | PA<br>ZT | -1.6 | -0.4 |

On average, a G-Z pair is less stable than an A-T pair by 0.27 kcal/mol, but the average masks a wide variation across stacks (standard deviation of ΔΔG° = 1.1 kcal/mol). Adjacent to A-T or other G-Z pairs, a G-Z pair substitution for an A-T pair is often (but not always) destabilizing. For example, a G-Z pair followed by an A-T costs 1.33 kcal/mol of stability relative to an A-T followed by an A-T. In some contexts, the G-Z substitution for an A-T pair is stabilizing, such as a G-Z following a P-Z, where the substitution is more stable than the A-T following a P-Z by −0.9 kcal/mol.

### Nearest Neighbor Stacking Enthalpy Changes

In addition to fitting free energy change stacking parameters, we also fit enthalpy and entropy change parameters using the same stacking model terms. These parameters enable the estimation of melting temperatures (T_m_s) for biotechnology applications and extrapolation of folding free energies to temperatures other than 37 °C.^25^ Like the free energy change parameters, the enthalpy and entropy change parameters were fit with a linear regression for P-Z pairs (Methods).

Table S6 shows the stacking enthalpy and entropy changes for stacks with P-Z pairs. The error estimates are larger for the enthalpy parameters than for the free energy change parameters as quantified by the fraction of the parameter value. This is expected; in fits to experimental data, the enthalpy and entropy changes are highly correlated and therefore free energy changes are determined with more precision.^22^

Table S7A provides the duplexes, the experimentally determined enthalpy changes, the fit enthalpy changes attributed to the P-Z, the fit values for comparison to experiment, and the residuals. Table S7B provides the equivalent information for the fit of the entropy changes. Given the larger average uncertainty in enthalpy and entropy changes from optical melting experiments and the lack of redundancy in experiments for G-Z pair stacks, we chose to not fit enthalpy or entropy change nearest neighbor parameters for G-Z pair stacks.

### Loop Stabilities

In addition to the helical stack parameters, the full set of nearest neighbor parameters required for secondary structure prediction includes parameters for estimating loop stabilities. To extrapolate a set of loop stabilities, we performed optical melting experiments on 32 duplexes that include dangling ends (Table S8), terminal mismatches (Table S9), single mismatches (from Wang et al., 2017; Table S10),^16^ and tandem mismatches (Table S10). 23 new optical experiments for loop systems were performed for this study; the other 9 had been reported previously.^16^ These four types of loops were chosen because sensitivity analyses for RNA secondary structure prediction identified these parameters as important for the precision of secondary structure prediction.^26, 27^

Dangling ends (single, unpaired nucleotides at the end of a helix) stabilize helix formation because the dangling nucleobase stacks on the terminal base pair.^28, 29^ For B-form DNA helices, 5′ dangling ends stabilize more than 3′ dangling ends. First, we studied the stability of 5′ dangling nucleotides on a P-Z pair (Table S11A and Figure 3A). On average, for dangling A, C, G, and T, the stability of the dangling end is 0.58 ± 0.07 kcal/mol less stable on a P-Z terminal pair as compared to a C-G terminal pair.

**Figure 3.**
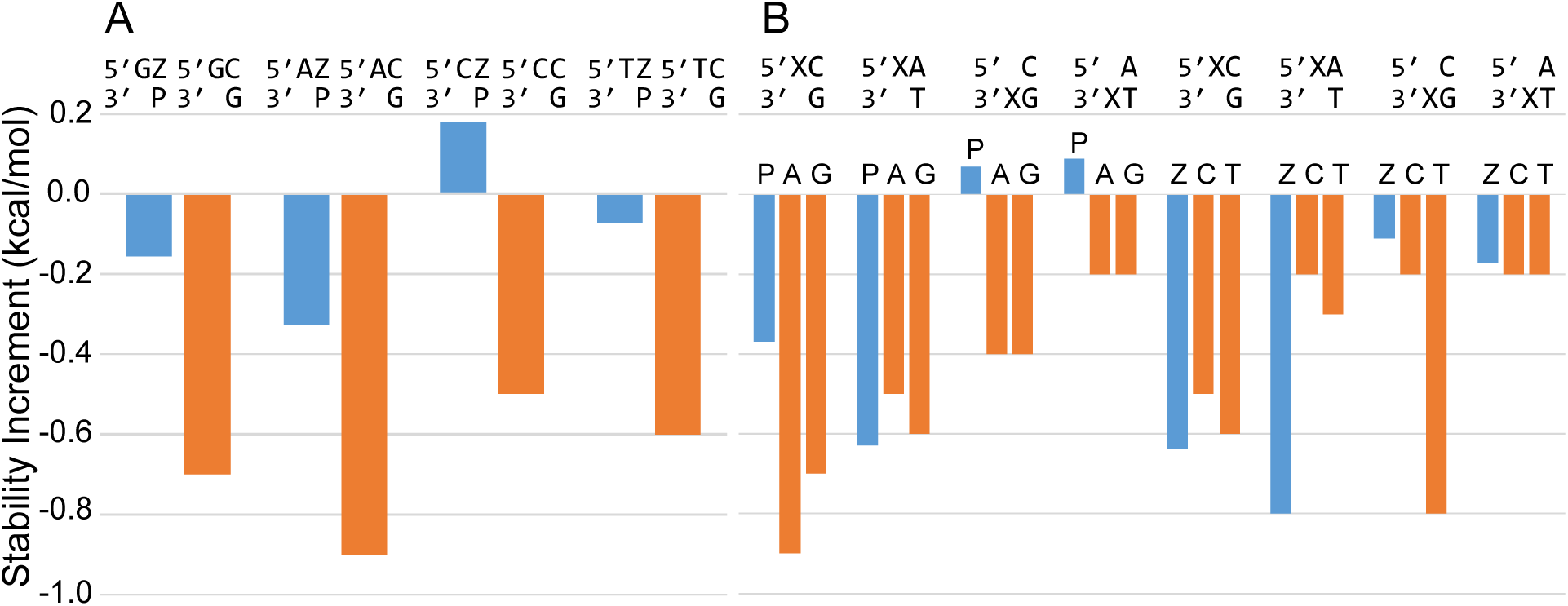
Dangling ends measured with P and Z nucleotides. Panel A. The stability of 5′ dangling nucleotides on Z-P pairs (blue), compared to C-G pairs (orange). The dangling end is less stabilizing on Z-P than on C-G pairs. Panel B. The stability of P and Z dangling ends, as compared to canonical purines and pyrimidines, respectively. The closing pair and orientation are indicated along the top. The dangling end nucleotide identity is directly above the blue bar.

We also studied the stability provided by P and Z dangling ends (Table S11B), which we found to be highly sequence dependent. Figure 3B shows the stability of P and Z dangling ends adjacent to Watson-Crick-Franklin pairs. A 5′ dangling Z adjacent to an A-T pair (−0.8 kcal/mol) is substantially more stable than either a C or T in the same context (−0.2 or −0.3 kcal/mol, respectively). In other cases, such as the 5′ dangling P (−0.37 kcal/mol), the stability is less than analogous A or G dangling bases (−0.9 or −0.7 kcal/mol). 3′ dangling P destabilized helix formation in the two contexts studied.

Terminal mismatches are single non-canonical pairs at the end of helices. These also stabilize helix formation because of stacking and hydrogen bonding. We studied the stability of P-P and Z-Z terminal mismatches adjacent to G-C and T-A terminal base pairs (Figure 4 and Table S12). The P-P mismatch next to a G-C pair is less stabilizing than the analogous purine mismatches G-G or A-A by 0.56 kcal/mol. The Z-Z mismatch next to a G-C pair, however, is similar in stability (at −0.76 kcal/mol) to the analogous pyrimidine mismatches C-C and T-T (with mean stability increment of −0.75 kcal/mol). The P-P mismatch next to a T-A pair is also similar in stability (−0.51 kcal/mol) to the analogous G-G and A-A mismatches (with mean stability increment of −0.5 kcal/mol). This contrasts to the Z-Z mismatch next to a T-A pair, which is more stabilizing than the analogous pyrimidine mismatches. The Z-Z mismatch next to T-A is −0.66 kcal/mol more stable (at −0.91 kcal/mol) than the mean of C-C and T-T next to T-A (at −0.25 kcal/mol). The general increased overall stability of dangling Z bases and terminal Z-Z mismatches is consistent with improved stacking of Z due to the nitro group.

**Figure 4.**
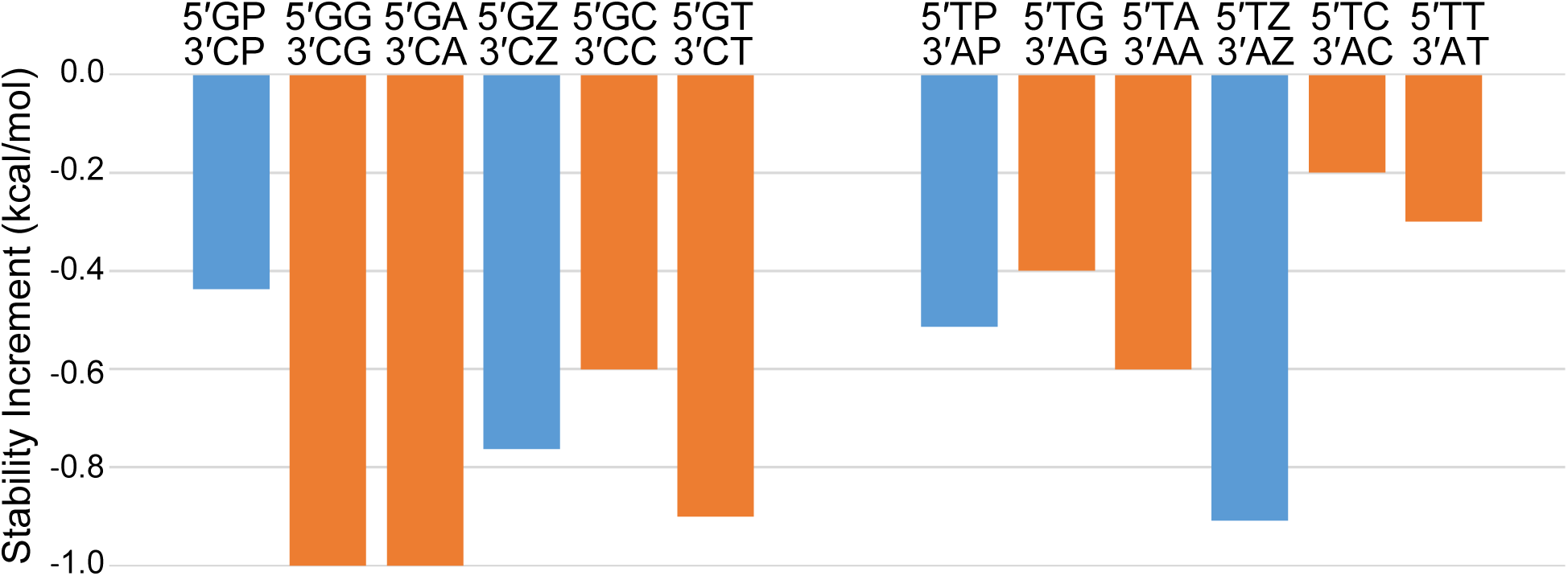
The stability of PP and ZZ terminal mismatches (blue), compared to GG, AA, CC, and TT terminal mismatches (orange). The left series have a G-C terminal pair and the right series have a T-A terminal pair.

We also measured terminal mismatches on terminal P-Z base pairs (Figure 5 and Table S13). Each of these terminal mismatches was less stable than the analogous mismatch on a terminal G-C pair. On average, this stability difference is 0.7 kcal/mol. We do not discern obvious patterns that explain the different results for these terminal mismatch stabilities. Given the conformational flexibility of ssDNA, there might be a variety of accessible structures for non-canonical pairing and stacking.

**Figure 5.**
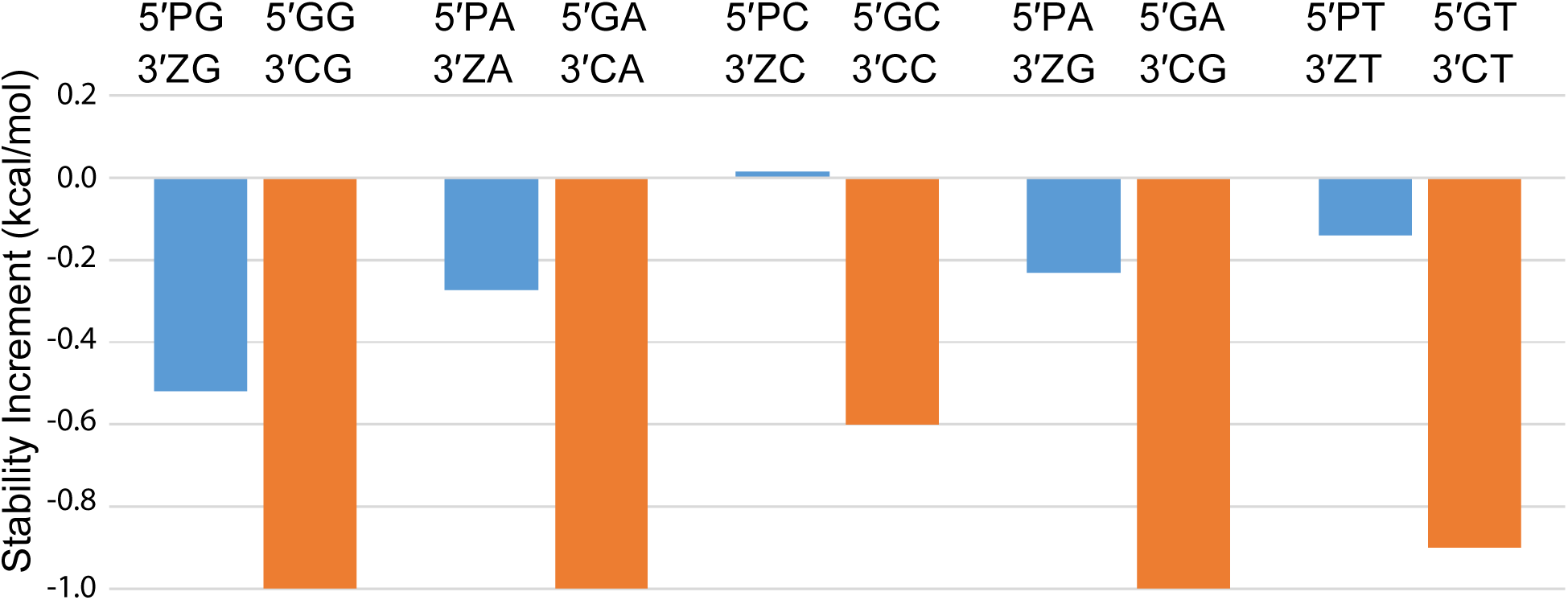
The stability increments of terminal mismatches on terminal P-Z pairs. The stabilities of analogous terminal mismatches on G-C pairs are shown.

We measured the stability of single mismatches (also called 1×1 internal loops): PC, PT, and ZA, by optical melting. In RNA structures, many non-canonical pairs that are capable of forming pairs with two hydrogen bonds have been well characterized.^30^ For the alphabet of nucleotides with P and Z, possible two-hydrogen bond non-canonical pairs of PC, PT, and ZA were previously proposed.^16^ There is a pronounced correlation of the stability with position in the helix, where single mismatches close to helix ends are stabilizing and single mismatches that are four base pairs separated from the helix end are destabilizing to duplex formation (Figure 6 and Table S14).

**Figure 6.**
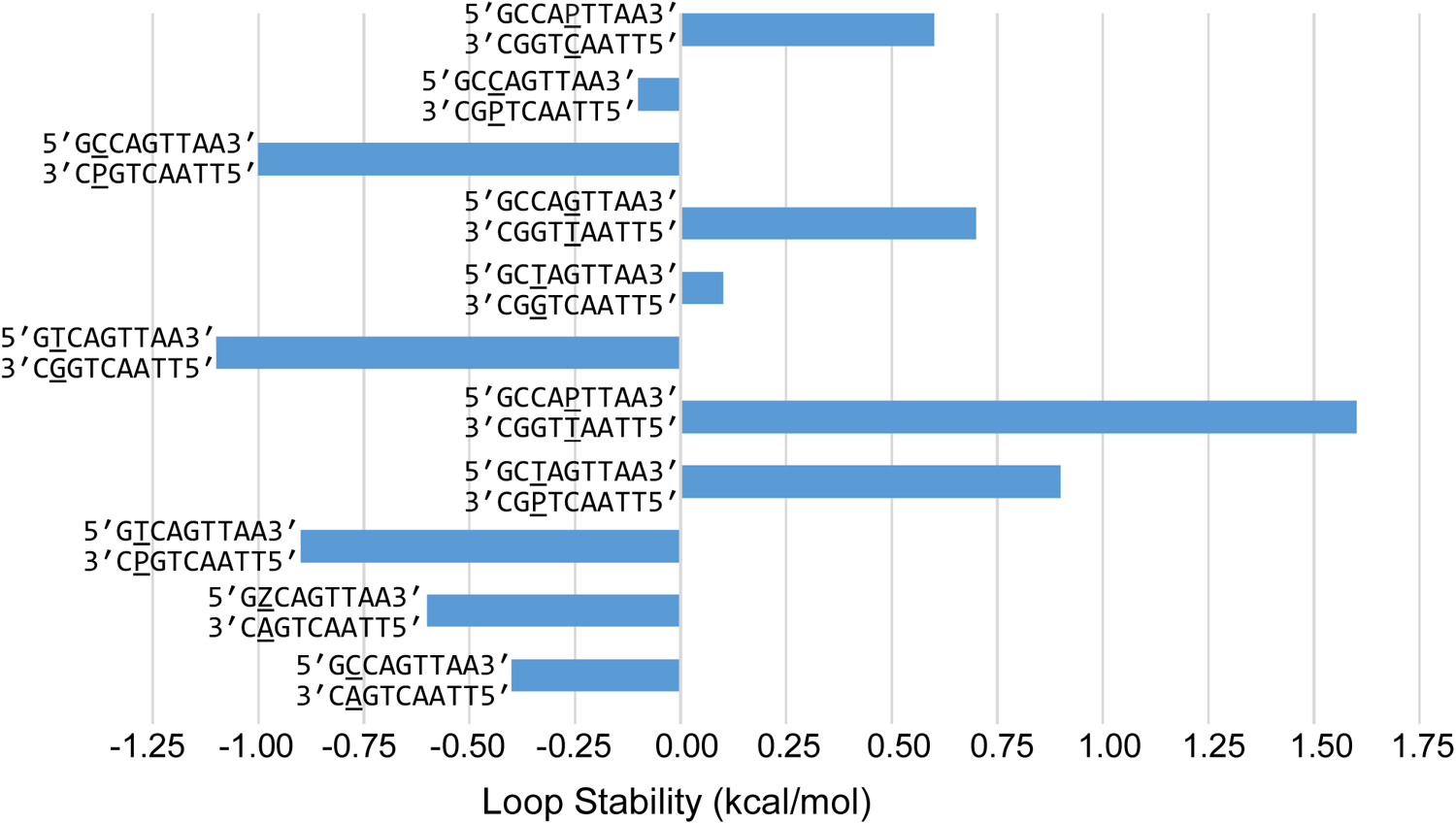
The stability of single mismatches (1×1 internal loops). Stability increments for the mismatch motif are shown, where the stabilities of the closing helices are subtracted from the duplex stability. The internal loops show a marked dependence on the distance from the helix end, where mismatches farther from helix ends are destabilizing and mismatches closer to helix ends are less destabilizing or stabilizing for helix formation.

We also observed this for control experiments with GT and CA mismatches. The existing database of optical melting experiments of model systems with natural DNA base mismatches focuses on mismatches with three canonical base pairs exterior to the duplex.^31–35^ Therefore, we developed single mismatch parameters for mismatches with P or Z by using the two total experiments with PC and PT mismatches in the center of the duplex.

We additionally measured the stability of two tandem mismatches (also called 2×2 internal loops) including a tandem Z-Z/Z-Z mismatch and a tandem P-P/P-P mismatch. Tandem mismatches have been studied extensively using canonical DNA nucleotides,^31, 36, 37^ and in one study were found to have a range of stabilities spanning 5.0 kcal/mol from stabilizing to destabilizing for duplex formation.^37^ We found that the Z-Z/Z-Z and P-P/P-P tandem mismatches are substantially more stable than analogous tandem pyrimidine or purine mismatches, respectively (Figure 7 and Table S15). On average, the tandem Z-Z/Z-Z and tandem P-P/P-P mismatch are −0.64 kcal/mol more stable than analogous tandem mismatches of canonical nucleotides. The stability of tandem Z-Z pairs might be explained by the Z-Z base pairing motif we characterized.^38^

**Figure 7.**
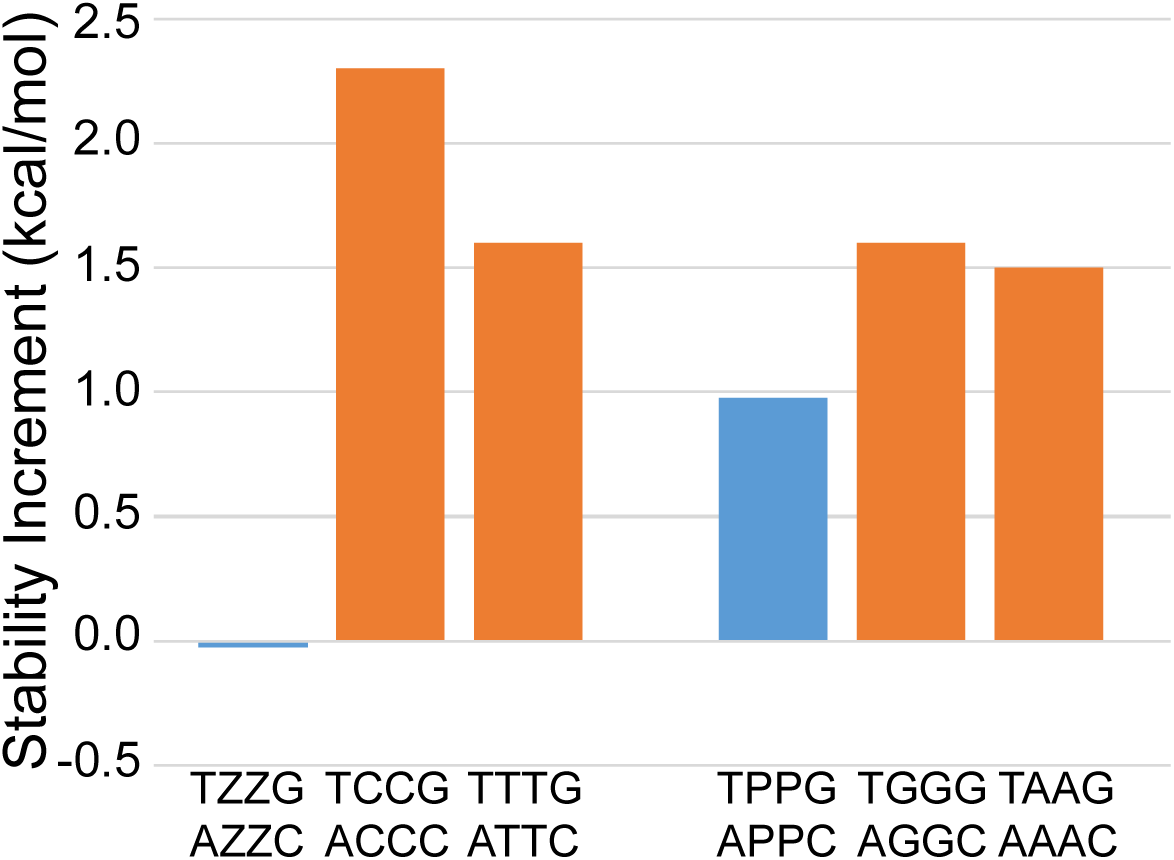
Stability increments for tandem mismatches (2×2 internal loops). The tandem ZZ and tandem PP mismatches are compared to tandem pyrimidine and tandem purine mismatches, respectively.

### Complete Nearest Neighbor Parameters

Using the optical melting stabilities, we extrapolated nearest neighbor terms for loops that contain P and Z or are closed by P-Z pairs. The details of the extrapolations are in the Materials and Methods section. RNAstructure was previously extended to accommodate extended alphabets beyond the canonical nucleotides.^18^ In summary, we developed a comprehensive set of thermodynamic parameters for A, C, G, and T, P, and Z, allowing C-G, A-T, P-Z, and G-Z base pairs, and we have integrated the expanded alphabet into RNAstructure such that any analysis that can be done for natural DNA can be done for the expanded alphabet.

### Tests of Sequence Designs that Include P and Z nucleotides

We tested the hypothesis that extension of the canonical base pairs with P-Z pairs would improve *in silico* designs of sequences that would fold into user-specified secondary structures. We used two types of structures for these tests: the Eterna 100 benchmark of structures,^19^ which is considered a challenge for automated sequence design, and a DNAzyme that had been discovered by *in vitro* evolution.^39^

To find sequences that fold to a specified secondary structure, we used the Design program in RNAstructure.^20^ The default parameters were used with one exception: we allowed isolated base pairs, i.e., base pairs that do not stack on adjacent base pairs in a helix, because the Eterna 100 structures contain isolated base pairs. Each structure was attempted five times (using the computer clock to seed the random number generator) and a maximum of five days for each of wall time was allowed using a single processor core. RNAstructure is capable of multithreading across CPU cores using OpenMP, but we used single threading to simplify the interpretation of the benchmarks. We considered success for a given target to be finding at least one sequence design that completed in the allowed five days.

Design was able to design sequences for 94 of the Eterna 100 structures using the canonical DNA bases only. When P-Z pairs were added to the design, 99 structures were successfully designed. When using P and Z, we only include P and Z in paired bases in the target structure. P-Z pairs are selected at a rate of 20%, G-C pairs are selected at a rate of 50%, and A-T pairs are selected with the remaining 30% (see Materials and Methods).

We then analyzed the results for the lowest NED for each alphabet of nucleotides. We focused on the lowest NED across calculations because Design uses stochastic refinement, and the NED of the final sequence therefore varies across calculations for the same structure.^20^ Figure 8 shows the lowest NED designed including P-Z pairs versus the lowest NED designed using only canonical nucleotides for the 94 test structures for which Design was able to solve both problems. The NED of sequences using P-Z pairs are lower in 89 cases, leaving only 5 cases where using P-Z is worse than canonical DNA nucleotides alone, and all of these 5 cases are in the lower left corner, where the NED using both alphabets is excellent (NED < 0.05).

**Figure 8.**
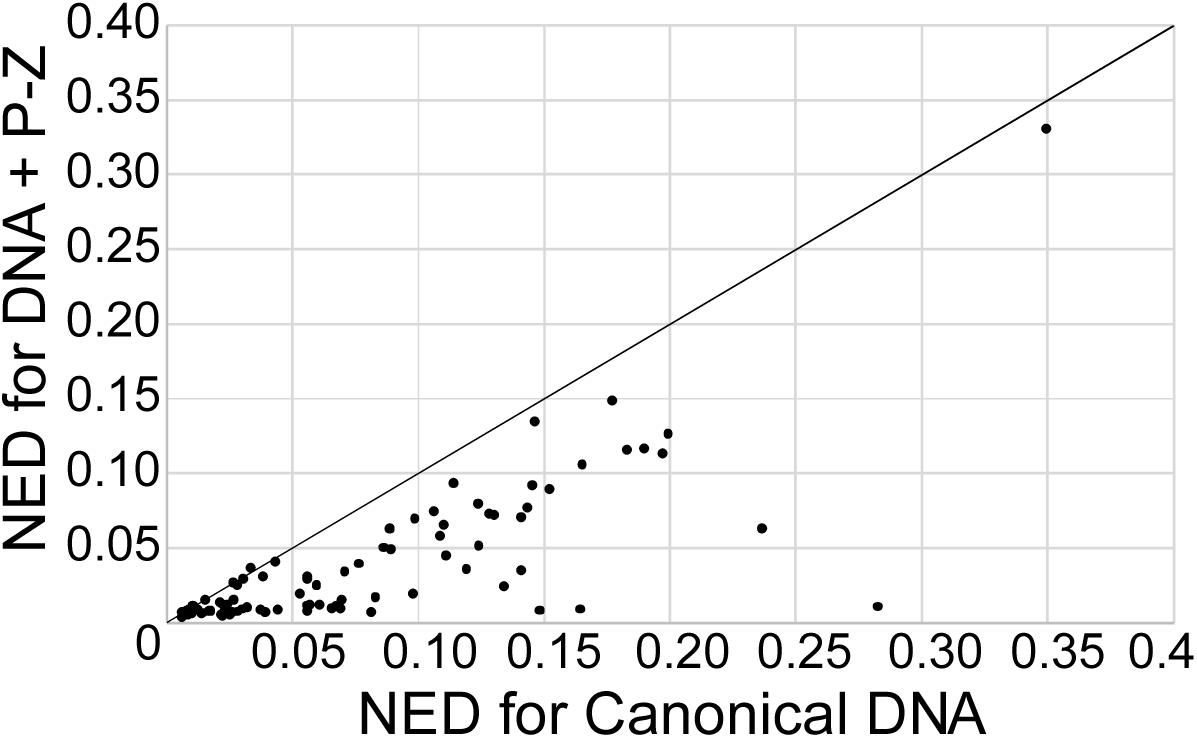
NED is significantly improved with the incorporation of P-Z pairs in designs (P=2.7×10^-13^). This is a plot of NED for Eterna 100 designs using DNA and P-Z pairs as a function of designs using canonical DNA only. The NED of the best of five calculations is shown. Each point is a single example problem from the Eterna 100 set. Points below the diagonal line (plotted as a visual guide) are cases where incorporation of P-Z pairs improved the designs.

A one-sided paired t-test indicated that the improvement in NED when incorporating P-Z pairs is significant (P=2.7×10^-13^). The average NED for the designs including P-Z is 0.036 and the average for NED for designs with canonical DNA is 0.073 for these 94 test cases solved with both alphabets. Figure S2 shows the analogous plot of the mean NED (rather than the minimum) using P-Z versus the mean NED using only canonical pairs. The same trend for mean is observed as with lowest NED, and the improvement in mean performance is again significant (P=1.0×10^-13^). We provide all the designed sequences in Supplementary Dataset 1, including the NED, the time used, and the random number seeds (to enable reproducibility).

We observed the most substantial improvement by incorporating P-Z pairs for the “Iron Cross” test (number 35), which improved in the best NED from 0.283 to 0.011. Figure 9 shows the target structure with the best DNA sequence and the best sequence for DNA with P-Z. The structures are color-annotated by the probabilities that nucleotides are in the desired structure. Like many of the other Eterna 100 test structures, this structure has prominent symmetry with four branches from a central multibranch loop, where each branch has a three-way multibranch loop with each helix of three base pairs. The canonical DNA design uses mostly G-C base pairs, but these can be promiscuous across the intended helices. The design that incorporates P-Z pairs has 23 P-Z pairs with the remaining 13 pairs as G-C. The target structure is stabilized by the incorporation of P-Z pairs, and alternative structures are preventable because there are additional possible combinations of P-Z and G-C pairs that form helices of three base pairs. Interestingly, P-Z pairs are chosen at a rate of 20% of pairs. The predominance of P-Z pairs after stochastic refinement in this case shows that P-Z are preferable in designed sequences that minimize NED.

**Figure 9.**
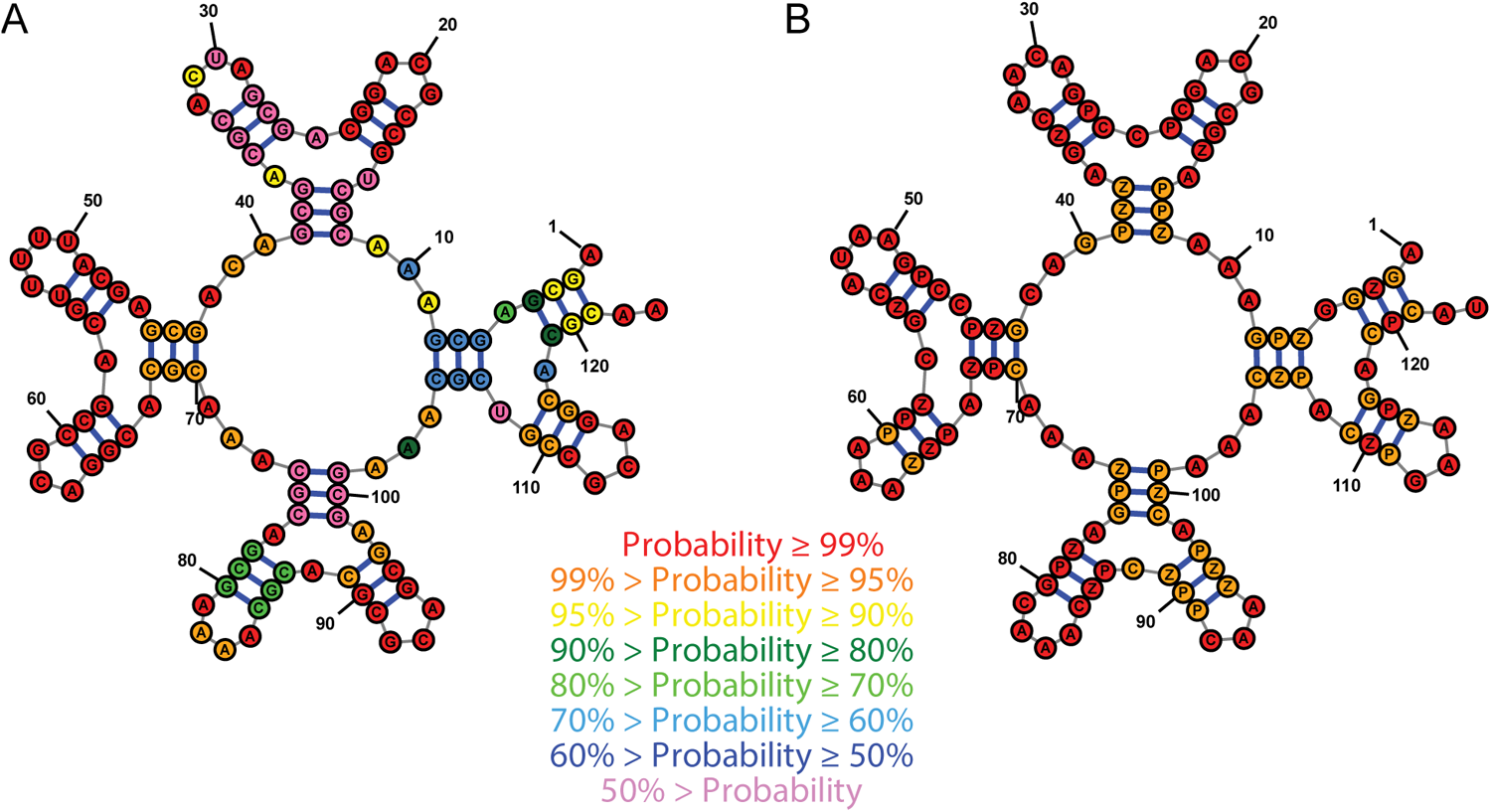
The incorporation of P-Z pairs improves the design of “Iron Cross,” problem 35 from the Eterna 100 set. Panel A shows the best design using canonical DNA only. Panel B shows the design incorporating P-Z into DNA. Bases are color-annotated with their probability of forming the correct structure, either the probability of folding into the specified target base pair or the probability of being unpaired in the target structure. The nucleotides in the P-Z containing sequence all form the desired structure with ≥ 95% probability. The structure composed of canonical DNA has a substantial number of nucleotides that are estimated to fold with < 50% probability to the target structure.

We previously noted that there is a tradeoff between time cost and NED quality when using stochastic refinement.^20^ In these calculations with Design, we used the default parameters, which limit the number of sequence refinements. In addition to observing an improvement in the quality of the designs (lower NED) when using P-Z pairs in the refinement, we also observe a decreased cost in average computational time. Figure S3 shows the average time across the five calculations that incorporate P-Z pairs as a function of average time across the five calculations that use only canonical DNA nucleotides for each Eterna 100 structure. In the calculation of average time, we use 5 days (the maximum allowed time) for calculations that failed to return a sequence. The use of P-Z nucleotides significantly improves the time cost (P=1.7×10^-3^).

In an attempt to solve additional Eterna problems, we extended the computational time allowed to ten days with Design. The one problem unsolvable (Gladius, structure 90) using P-Z remained unsolved. Three additional problems, however, were solved by extending the canonical DNA designs to 10 days (Supplementary Dataset).

99 of the Eterna 100 benchmark structures have Eterna player solutions provided, with two RNA sequences being provided for each. We compared the NED of our designs including P-Z pairs to the Eterna solutions. Eterna designs were originally evaluated computationally as to whether the sequence is predicted to fold with lowest free energy to the target structure. This can be achieved whether or not the NED is relatively low,^13, 40^ and therefore we are evaluating the sequences with a different metric than for which they were designed. We compared the lowest NED for our solutions using P-Z to the lowest NED using Eterna player RNA sequences (Supplemental Fig. S4). Our designs, including P-Z, have significantly lower NED than the Eterna RNA designs (P=4.28×10^-19^).

We additionally tested including P-Z base pairs in designing DNA with the same secondary structure as a DNAzyme:substrate complex, a unimolecular version of the CT10.3.29.M5 sequence bound to its target.^39^ The catalytic core of the DNAzyme, which includes several conserved or essential nucleotides, was specified as unpaired; the current implementation of the Design algorithm does not allow the sequence of to be constrained. We designed 10 sequences including P-Z pairs and 10 sequences using only canonical DNA nucleotides. Table S16 reports the results in terms of NED and time. The NEDs for the DNAzyme designs were excellent (i.e. low) for either alphabet. Using P-Z pairs the lowest NED solution was 0.031, and using canonical DNA the best NED solution was 0.041. The average time performance using P-Z pairs was better and more consistent than when only using canonical nucleotides. The mean time with P-Z pairs was 1594 ± 134 s and the mean time for canonical nucleotides was 3232 ± 1770 s. The improvements when using P-Z pairs are significant for NED (P=1.90×10^-3^) and time (P=8.46×10^-3^). Within each set of designs, there was no correlation between the time required and the quality of the design.

## DISCUSSION

In this contribution, we tested the hypothesis that synthetic base pairs can improve the quality of DNA nanostructure designs. We used optical melting to study the folding stability of DNA containing P and Z Hachimoji nucleotides. These data informed a new nearest neighbor model for folding, including P-Z pairs, G-Z wobble pairs, and loops containing P or Z. Using the Design program in RNAstructure,^20, 41^ which was recently enhanced to allow prediction with nucleotides alphabets beyond the canonical nucleotides,^18^ we demonstrated significant improvements for *in silico* sequence design quality.

Prior work studying RNA nearest neighbor parameters demonstrated that a subset of parameters needs to be determined with high precision to precisely estimate base pairing probabilities. That work focused our efforts here to measure folding stability for model systems that could inform those important parameters. This approach was recently validated for the derivation of folding free energy parameters for RNA including m^6^A.^18^ A set of 98 duplexes with or without N^6^-methylation, including helices, bulge loops, internal loops, dangling ends, and terminal mismatches, was studied and this demonstrated that the parameters extrapolated for m^6^A were approximately as accurate for estimating folding stability as those for canonical bases only.^42^

There are limitations to accuracy of our parameters. For example, single mismatch parameters for canonical DNA do not account for the positional dependence of the mismatch stability.^29^ This is because the current experimental database of DNA mismatches is limited to mostly mismatches distant from helix ends and because current structure prediction packages are not capable of accounting for this dependence. For single mismatches containing either P or Z, our parameters are based on the most destabilizing values that are consistent with being away from helix ends. This is currently a limitation in parameterization for both DNA and DNA+PZ folding alphabets. Both DNA parameters and DNA+PZ parameters would benefit from additional experiments to guide the determination of helix-position-dependent rules.

Comparison of the refined nearest neighbor thermodynamic parameters obtained here with previous DNA and PZ parameters (Figure 10) confirms that P-Z pairs are generally more stable than natural sequence DNA. Our revised parameters for PZ-containing dinucleotides generally show more negative ΔG°_37_ for base pair formation than the Hachimoji set, but the differences are small. As described above, one significant difference is that we do not include a free energy penalty for a terminal P-Z pair. The free energies for the GZ pairs are more variable, with the most stable G-Z containing stack, PG/ZZ, being slightly more stable than PP/ZZ, while the least stable GZ/ZG stack is substantially destabilizing. This is not due solely to having two mismatches in the stack, as ZG/GZ is quite stable.

**Figure 10.**
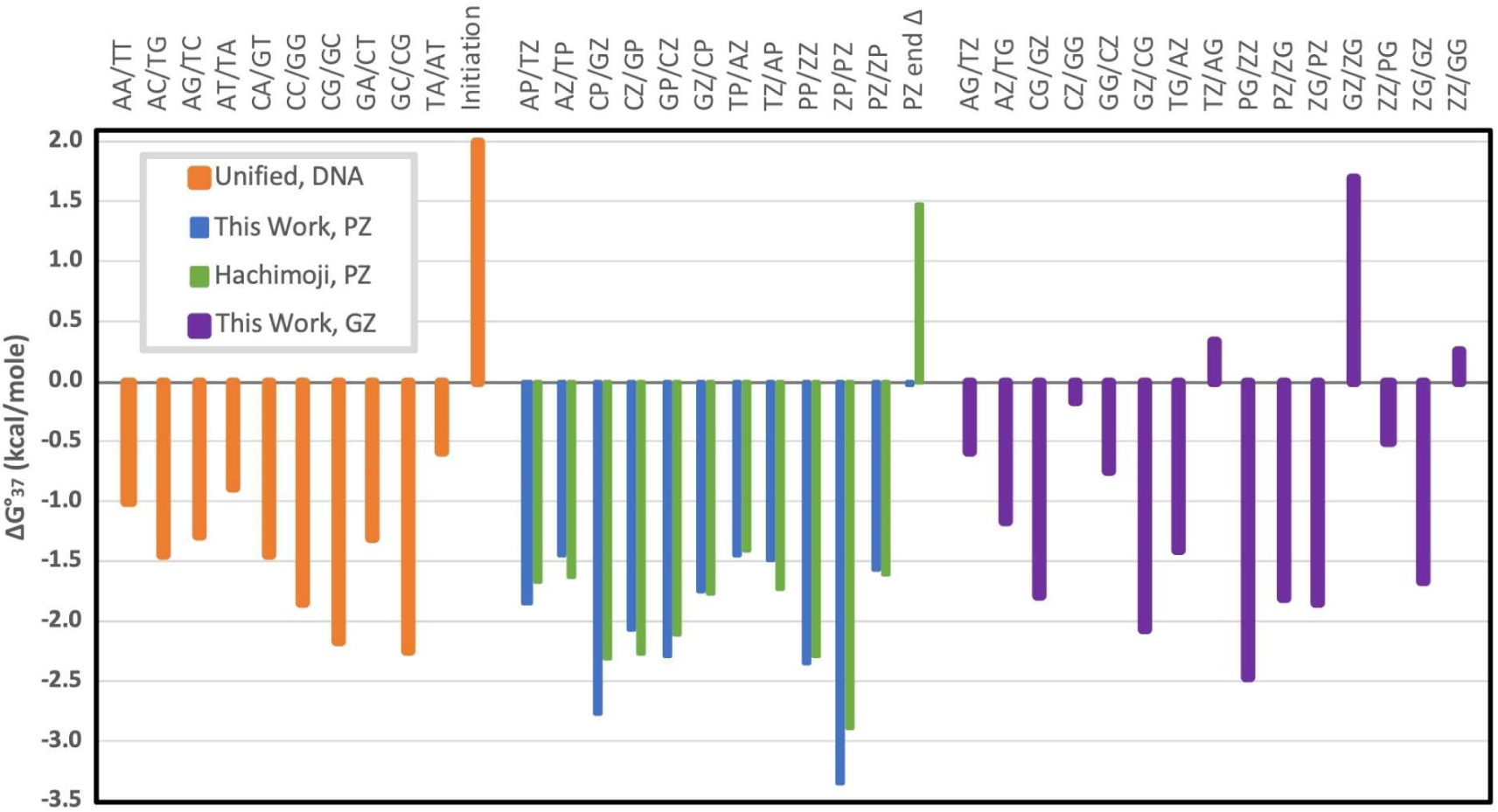
Nearest neighbor free energy parameters for stacks containing natural DNA, P-Z pairs, and the stable G-Z mismatch.

There is a notable effect of the position of the Z base. For the P-Z data sets, stacks with Z at the 5′ side (e.g. XP/YZ, including ZP/PZ and PP/ZZ) have an average ΔG°_37_ = –2.34 kcal/mol whereas stacks with Z at the 3′ side (XZ/YP, including PP/ZZ and PZ/ZP) have ΔG° = –1.78 kcal/mol. For G-Z pairs, the average ΔG°_37_ for 5′ Z is –1.19 kcal/mole and for 3′ Z it is –0.65 kcal/mole: the absolute difference is very similar to P-Z but the relative difference is much larger. Examination of X-ray crystal structures (PDB accessions: B-form 4XO0, 6MIG, and 6MIH and A-form 4XNO) does not suggest an obvious reason for this dramatic preference.^43^ One might guess that the potential stacking of the Z-NO_2_ group could be particularly stabilizing, but the Z base appears to stack much better on its 5′ neighbor than on its 3′ neighbor. The 5′ Z is solvent exposed, and it is surrounded by ordered water in 4XNO.

Our results highlight a case where the use of P-Z pairs is crucial for high quality (i.e. low NED) designs. Structures with extensive symmetry (such as the “Iron Cross” example in Figure 9) benefit from the additional space of sequence options that are available when P-Z pairs are included. In the absence of P-Z pairs, the three base pair helices, which need the stabilization of G-C pairs, exhaust the possible sequences and then alternative structures form when the helix sequences pair incorrectly. We hypothesize that this important diversification of sequence effect plays a smaller, but beneficial, role in other sequence designs that are not as symmetrical as “Iron Cross.”

Figure 8 demonstrates cases when the NED of the best sequence designed including P-Z pairs is worse than the best sequence designed with DNA only. Given enough search time or enough search attempts, this would not happen because the DNA designs including P-Z pairs could find the better DNA-only solution. We observe these cases because Design stochastically refines the sequences and returns solutions when it has achieved a specified threshold NED or when it has exhausted a specified number of trials. For this work, we chose to not refine the default parameters for the search, although it might be possible to improve the best NED by changing these settings.

As alphabets expand, the number of parameters needed to describe even the simplest nearest-neighbor models expands combinatorially. Comprehensive description of mismatch and internal loop thermodynamics is out of reach for low-throughput measurements like those described here. Systematic evaluation of which parameters are most critical,^26, 27^ high throughput measurements of enthalpy and entropy as well as free energy changes,^44^ position-dependent rules, and eventually integration of 3-D motif stability^45, 46^ will be necessary to tackle more demanding design problems like structures that change their folds upon ligand binding or as conditions change.

## METHODS

### Oligonucleotide Synthesis, DNA Melting, and Analysis of Melting Curves

P/Z containing DNA oligonucleotides were synthesized as described,^16^ and unmodified oligonucleotides were purchased from IDT (Coralville, Iowa). The purity of all oligonucleotides was evaluated by denaturing PAGE, staining with SYBR Gold, and imaging at 312 nm. We note that Z-containing oligonucleotides quench SYBR fluorescence in the gel; Z has a broad absorption peak centered at about 380 nm.

Absorbance melting curves were performed as described for the P-Z pairs and loop energy oligos.^16^ For G-Z melts we improved the standard procedure in several ways. We acquired absorbance vs. temperature curves for each of the single strands individually, both to determine their diluted stock concentrations directly and to use actual data instead of a linearly extrapolated ssDNA trendline. We carried out simultaneous dsDNA melts at two concentrations, nominally 1 µM and 5 µM total strand concentration, C_T_, which enabled rapid identification of pipetting errors and set constraints on the concentrations of the two individual strands in the dsDNA melt cuvettes. We used equations for predicting total absorbance that explicitly consider unequal concentrations of the single strands and the absorbance of the remaining excess single strand.

We carried out global fits to both of the absorbance vs temperature curves with a set of six fit parameters: two ssDNA concentrations for each melt (the other two being determined by the total absorbance), a common % hypochromicity for the duplex, a common slope for the dsDNA absorbance vs T, ΔH°, and ΔS°. The fit was repeated four times to consider each pair of possibilities in each melt for which strand is in excess. The fit with the lowest residual that also had minimal errors in concentrations relative to nominal values was selected. This procedure allows explicit correction for pipetting errors, which are typically in the range of 5%, and the global fitting to dsDNA absorbance decreases the total number of parameters by two for each melt. We observe that typical estimated errors in ΔH° and ΔS° are reduced from 8-10% to about 5% by following this procedure. It could readily be extended to melts at additional C_T_, as is done with the alternative ln(C_T_) vs 1/T_m_ procedure for determining ΔH° and ΔS°.^47^ An example of the output from our Matlab routines is provided in the Supplemental and the code is available at GitHub at https://github.com/jasondkahn. We note that other recent work is available for improved fitting of optical melting data.^48^

### P-Z and G-Z Stacking Free Energy Change Nearest Neighbor Fits

A custom Python program was written to fit the P-Z nearest neighbor parameters. First, the sum of the Watson-Crick-Franklin stacks, the intermolecular initiation, and the symmetry correction (if needed for self-complementary duplexes) was subtracted for each duplex. For compatibility with RNAstructure, we used the parameters that are provided with the RNAstructure package.^41^ Then, the nearest neighbor stacks were fit by linear regression to the remaining stabilities with the statsmodels ordinary least-squares class (OLS).^49^ This fixes the stability of the Watson-Crick-Franklin stacks with their existing values. Similarly, the stacks with G-Z pairs were fit with a separate custom Python program, fixing the values of Watson-Crick-Franklin and P-Z stacks. Error estimates for parameters are the standard errors of the regression.

### P-Z and G-Z Stacking Enthalpy and Entropy Change Nearest Neighbor Fits

To fit enthalpy change nearest neighbor parameters, a custom Python program was written. The Watson-Crick-Franklin stack enthalpy changes and intermolecular initiation enthalpy changes were subtracted from the experimentally determined enthalpy for each duplex. The Watson-Crick-Franklin enthalpy change parameters were those reported by SantaLucia and Hicks.^29^ Then the enthalpy change nearest neighbor parameters were fit by linear regression using the statsmodels ordinary least-squares class (OLS).^49^ A second custom Python program was used to fit the G-Z enthalpy change stacking parameters.

The enthalpy changes of the Watson-Crick-Franklin and P-Z stacking enthalpy changes and initiation were subtracted from the total experimental-determined enthalpy changes. The G-Z enthalpy change terms were then fit to this remaining stability by linear regression. Error estimates for the enthalpy change parameters are standard errors of the regression. Following the same methods used to fit the enthalpy change nearest neighbor parameters, two additional Python programs were written to fit the entropy change nearest neighbor parameters for stacks with P-Z pairs and for stacks with G-Z pairs.

### Determination of Loop Motif Stability Increments

Loop stability increments are determined by subtracting helix stabilities from the optical melting stability determined for model systems. In this work, non-self-complementary duplexes were used to determine dangling end, terminal mismatch, and internal loops. For the dangling ends and terminal mismatches, the stability increment of the motif is determined by subtracting a reference helix stability from the stability of the duplex with the motif:

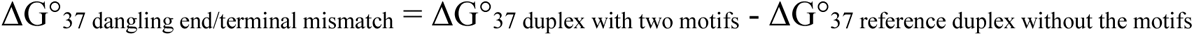

The stability of the reference duplex is estimated with stacking nearest neighbor parameters.

For internal loops, the stability is the total stability of the duplex minus the helical stacks (estimated with nearest neighbor parameters):

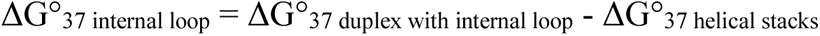

### Extrapolation of Loop Nearest Neighbor Parameters

For the basis of our free energy change nearest neighbor parameters, we used the DNA parameters incorporated in RNAstructure. For the DNA+PZ alphabet parameters, we removed GT pairs, which were traditionally included as part of DNA secondary structures. GT pairs are less stable than AT pairs and GZ pairs,^31^ and they are treated as mismatches in the DNA+PZ alphabet.

For 5′ dangling ends on terminal Z-P pairs, we used the experimental values (Table S11A) for A, C, G, and T. The dangling end parameters for DNA are not very sequence dependent. The largest differences are for 3′ dangling ends on terminal CG pairs, which range from −0.4 to −1.1 kcal/mol in stability. For other terminal pairs, the dangling end stabilities (3′ or 5′) are in a range of 0.4 kcal/mol.

Therefore, we approximate a 5′ dangling A, C, G, or T on a terminal P-Z pair as the mean of the measured values on the Z-P pair (−0.1±0.2 kcal/mol). For 3′ dangling ends, we are conservative and use the same −0.1 kcal/mol. For dangling ends on P-Z or Z-P terminal pairs, we use the experimental values when available (Table S11B). For 3′ P dangling ends that were not measured, we use the mean value of those measured (Table S11B; 0.1 kcal/mol). For 5′ P dangling ends that were not measured, we use the mean value of those measured (Table S11B; - 0.5 kcal/mol). Similarly, for 3′ or 5′ Z dangling ends that were not measured, we use the mean of the measured values, which is −0.1 or −0.7 kcal/mol, respectively (Table S11B).

For terminal mismatches, we used our measured values when those were available. For PP terminal mismatches that were not measured, we used the mean from other PP mismatches (Table S12; −0.5 kcal/mol). Likewise, for ZZ terminal mismatches not measured, we used the mean from other ZZ mismatches (Table S12; −0.8 kcal/mol). For terminal mismatches on P-Z, Z-G, or G-Z pairs, including mismatches with P or Z (but not PP or ZZ), we use the mean value of the terminal mismatches on Z-P pairs (Table S13; −0.2±0.2 kcal/mol). Terminal mismatch parameters are used in secondary structure prediction for mismatches in exterior loops, hairpin loops, internal loops (other than 1×1, 1×2, 2×2, or 1×n, where n>2), and multibranch loops. The terminal stack parameters also provide one of the terms for the stability of mismatch-mediated in coaxial stacking.^50–52^

For single mismatches, a.k.a. 1×1 internal loops, parameters were informed by experiments reported in Table S14. We use mismatches in the center of adjacent canonical helices to be consistent with prior studies of DNA single mismatches. The single mismatch parameter table in RNAstructure contains all sequences (including the mismatch and two closing base pairs). When an experimental value has been measured, we use that value. The following extrapolations were used to determine parameters involving P or Z: We map PZ closure to entries for GC closure and we map GZ closure to entries for AT. In the loops, we map Z to C and P to G. If the mapping results in a Watson-Crick-Franklin pair, we remap to an AC mismatch, preserving the purine-pyrimidine orientation. We then lookup the entry as mapped to canonical nucleotides and stabilize mismatches with P or Z by −0.6 kcal/mol, which is the mean additional stabilization observed for the measured mismatches (Table S14) as compared to analogous AC mismatches.

For tandem mismatches, we use a similar approach as single mismatches. RNAstructure uses a table with all possible sequence entries. We use the same remapping used for single mismatches to look up an entry composed of canonical nucleotides. Then, for each mismatch with a P or Z, we stabilize by an addition −0.6 kcal/mol, which is the mean stabilization per mismatch observed for tandem mismatches as compared to analogous loops with canonical nucleotides (Table S15).

For 2×1 internal loops, RNAstructure also uses a table with all sequence entries. Similar to single mismatches, we map entries with P or Z back to entries with canonical nucleotides only. For loops that contain either P or Z, we stabilize the loop by −0.6 kcal/mol, the average stabilization observed per mismatch with P or Z in single mismatches (Table S14).

Following the nearest neighbor parameters for DNA, the PZ alphabet tables for first mismatches in hairpin loops, mismatches at helix ends in internal loops (larger than 2×2 internal loops), and for mismatches at helix ends in coaxial stacks are set equal to the terminal mismatch values.

### Design Calculations

The Design program from RNAstructure was used for the calculations.^20, 41^ We modified the program to allow isolated base pairs, i.e. base pairs that are not stacked on adjacent base pairs, and this revision will be available in RNAstructure version 6.5. The other parameters were set to defaults. The software was compiled using GCC version 4.8.5. All calculations were run on one core on machines equipped with two Intel Xeon CPU E52695v4 (2.10 GHz) processors and 124 GB of RAM. The CPU time was extracted from a Slurm queuing report.

Design calculations were all run using the thermodynamic parameters that include P and Z nucleotides. Design randomly selected pairs and unpaired nucleotides for initial sequences and then subsequently for the refinement of the sequence. We extended Design to allow user-specified biases for the selection of pairs and unpaired nucleotides. These biases are used to initialize the sequence and also in the selection of revised sequences during the iterative refinement stage. For calculations with only canonical nucleotides, G-C and A-T pairs were selected with equal probability. For designs that included P-Z pairs, P-Z was selected with probability of 20%, G-C with a probability of 50%, and A-T with a probability of 30%. For loop sequences, A, C, G, and T were selected with probabilities of 60%, 10%, 10%, and 20%, respectively. This reflects a bias for A occurring in loops.^53^ P and Z were not selected for incorporation in loops.

### Statistical Tests

For analysis of the Eterna 100 set designs, we used SciPy 1.7.3 to perform paired t-tests for NED or time.^54^ Our null hypothesis is that the quality of designs with PZ pairs are not better (NED or time) than the designs with canonical nucleotides only. Similarly, for the DNAzyme calculations, our null hypothesis is that the quality of designs with PZ pairs are not better (NED or time) than the designs with canonical nucleotides only. We tested this null hypothesis using Welch’s t-tests for NED or time. For all tests, we used a one-sided test with the type I error rate, α, set to 0.05.

## Author Contributions

T.M.P. and M.Z. carried out folding and design calculations. H.S. and D.H.M. performed regression analysis. T.M., K.K.S., and X.W. performed absorbance melting experiments. S.H. synthesized, purified, and characterized all of the oligonucleotides containing non-standard nucleotides. R.J.P and S.A.B consulted on all aspects of the work, and reviewed the manuscript. J.D.K. wrote analysis programs and supervised experimental work. D.H.M. supervised the computational work. T.M.P., J.D.K, and D.H.M. wrote the manuscript with input from all of the authors.

## Funding

This work was supported by NIH grant R35GM145283 to D.H.M. and an SBIR award from the U.S. Army (W911NF-12-C-0060) to Celadon Laboratories (later DNA Analytics). This work was also supported by R01GM141391-01A1 to S.A.B.

## Notes

DNA Analytics applied for a U.S. patent, PCT/US23/64288 (Generating Parameters to Predict Hybridization Strength of Nucleic Acid Sequences), to protect the applicable optical melt curve fit methods.

## ACKNOWLEDGEMENTS

Sequence design calculations were performed using resources provided by the University of Rochester Center for Integrated Research Computing.

## Abbreviations

e.u.: entropy units (cal mol^-1^ K^-1^)
NED: normalized ensemble defect

---

For Table of Contents Only:

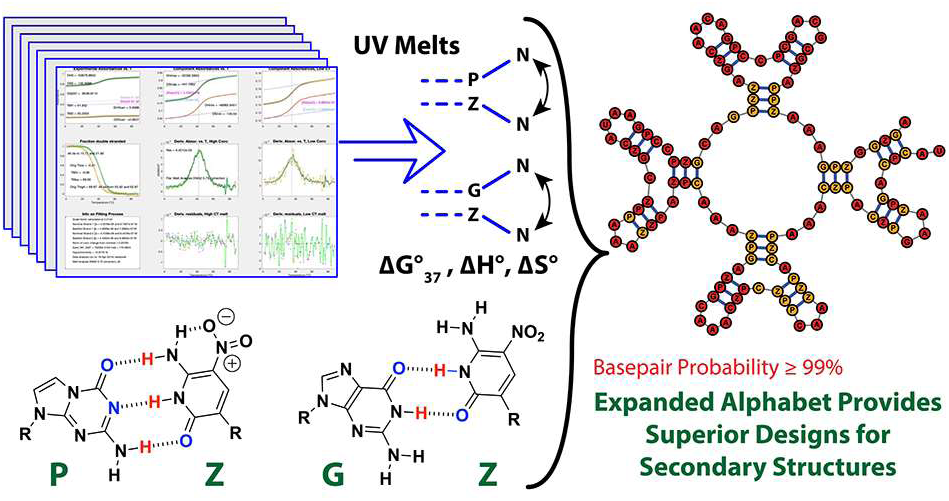

