## Supplemental Tables and Figures for "DNA Structure Design Is Improved Using an Artificially Expanded Alphabet of Base Pairs Including Loop and Mismatch Thermodynamic Parameters"

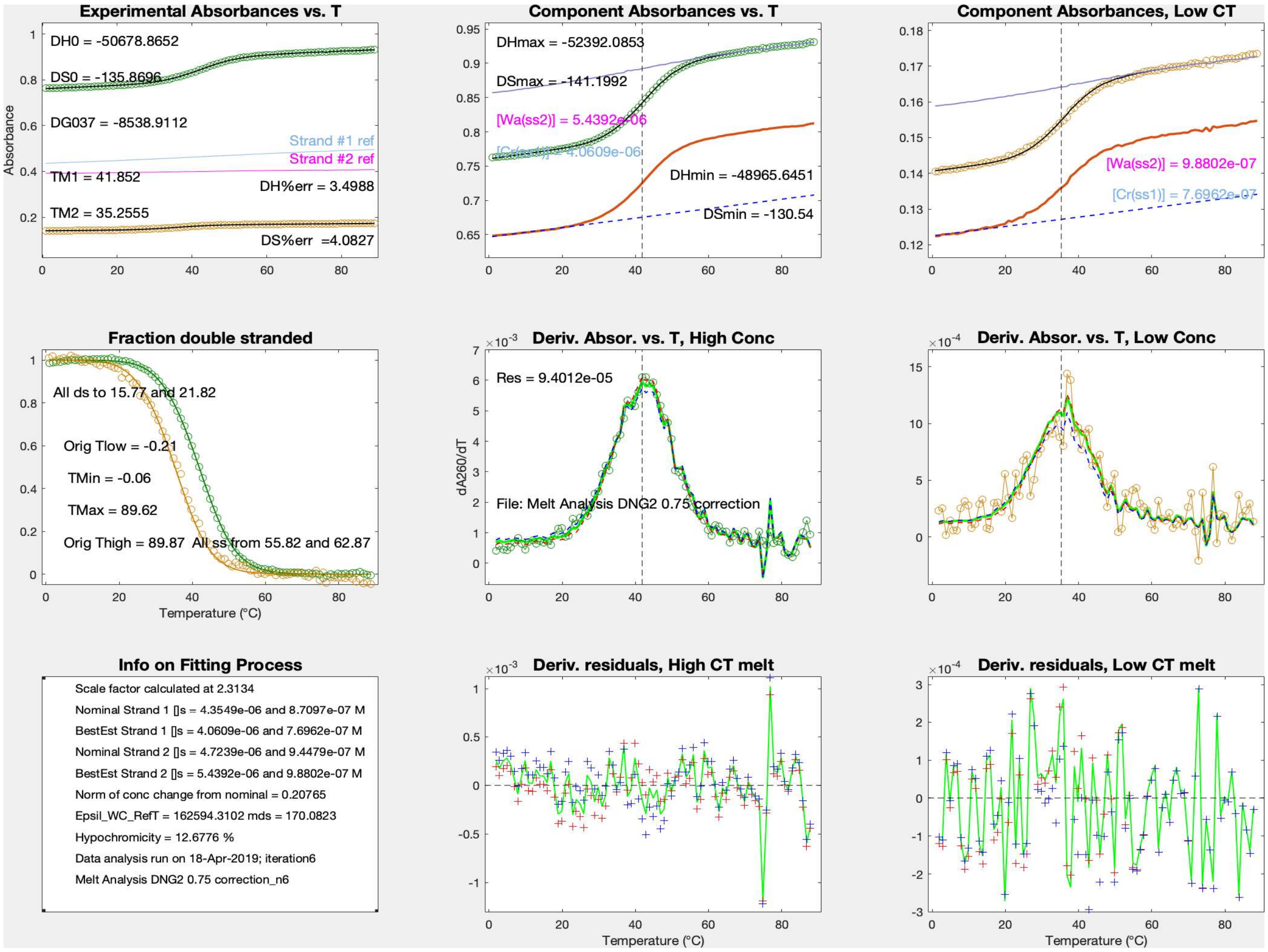


Figure S1. Sample output from global fitting to two optical melting experiments. The plots show the raw data, derivative plots, and analysis for two concentrations. The Wa strand is in excess by definition. Nominal concentrations of the working stocks are determined from strand #1 and strand #2 reference curves in the top left plot. The lower orange curves in the two plots at the top right show the experimental curve minus the absorbance due to the excess single strand. Note that the y-axis does not start at zero. Also, note that because of LeChatelier's principle, the orange curve would not actually be observed for equal single strand concentrations. In the algorithm, the global fit to normalized absorbance change weights the higher-concentration curve more heavily, according to the square root of total absorbance. The Matlab routines generate four analogous plots for the four possible combinations of which strand is in excess in each melt.

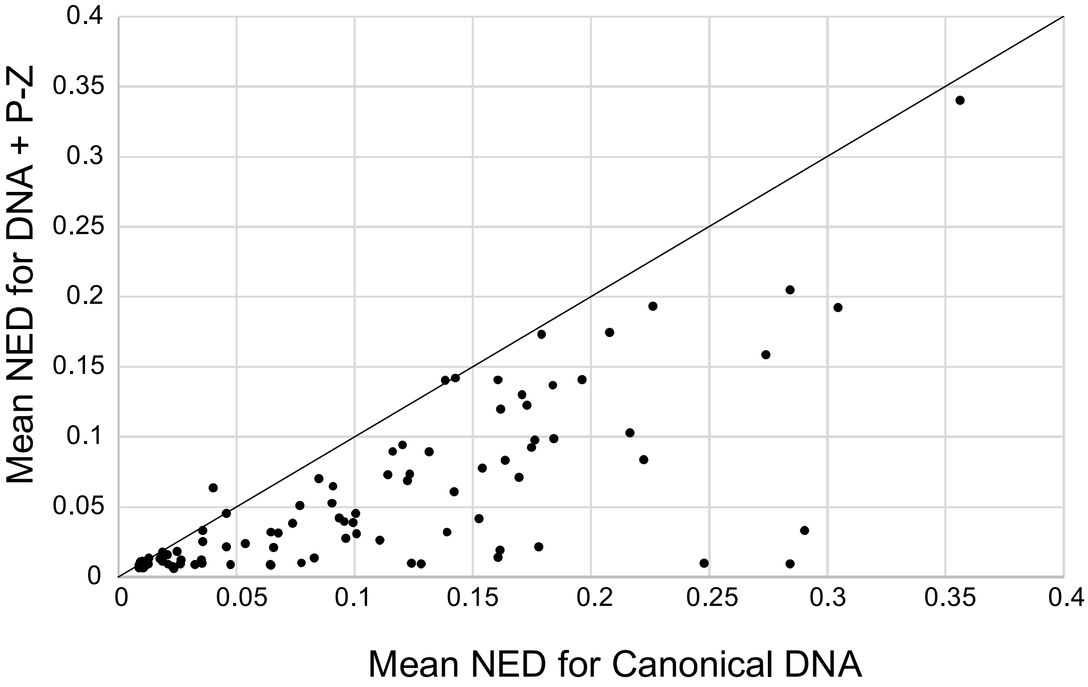


Figure S2. Mean NED is significantly improved with the incorporation of P-Z in designs ($P=1.0\times10^{-13}$). The mean NED for designed sequences that include P-Z pairs as a function of NED for designed sequences that use only canonical DNA nucleotides. Points below the diagonal (shown for reference) are test examples with improved (lower) NED when P-Z pairs are used.

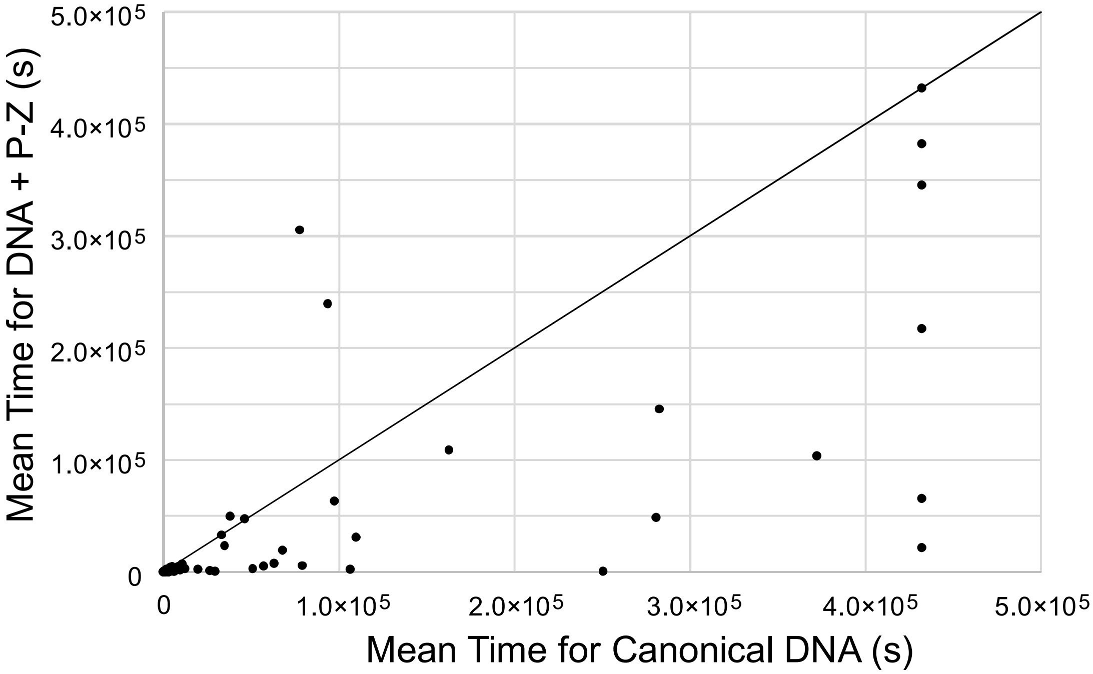


Figure S3. Average time is significantly improved with the use of P-Z pairs as compared to using only canonical DNA nucleotides ($P=1.7\times10^{-3}$). The mean time for five Design calculations with P-Z pairs is plotted as a function of mean time for five Design calculations using only canonical DNA nucleotides. Each point is a single problem from the Eterna 100 set. Failed Designs are those that reached the 5 day limit (432,000 seconds). Six problems were not solved in the five attempts using DNA only, and these are points that align vertically at 432,000 s. One problem was solved with neither DNA nor with DNA including PZ and this point is on the diagonal at 432,000 s. 70 problems were quick (less than 3 hours or 10,800 seconds) to solve for both P-Z and canonical DNA only. These points are clustered in the lower left of the plot.

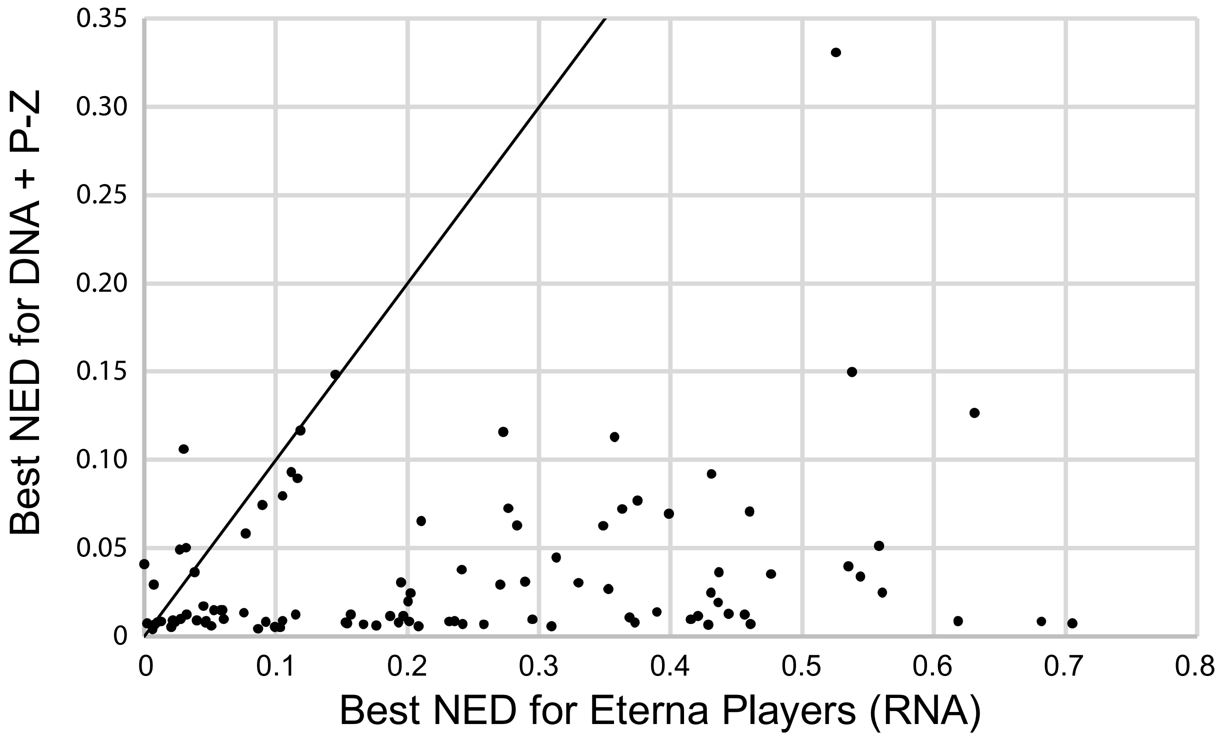


Figure S4. The best NED designed for DNA with P-Z pairs is significantly better than the best NED designed for RNA by Eterna Players ($4.28\times10^{-19}$). The diagonal is shown as a visual aid. Solutions below the diagonal have lower (better) NED for RNAstructure Design than for Eterna players.

Table S1. Optical melting data for 13 new duplexes designed to determine P-Z stack nearest neighbor parameters.

| Sequence 1 5′->3′ | Sequence 2 3′->5′ | ΔH° (kcal/mol) | ΔS° (cal mol$^{-1}$ K$^{-1}$) | ΔG°$_{37}$ (kcal/mol) |
|---|---|---|---|---|
| GCACAGATC**P** | CGTGTCTAG**Z** | -75.4 ± 2.9 | -205.3 ± 9.0 | -11.71 ± 0.15 |
| GCACAGTTT**Z** | CGTGTCAAA**P** | -72.4 ± 2.4 | -199.4 ± 7.5 | -10.61 ± 0.10 |
| GCACA**P**CTGA | CGTGT**Z**GACT | -72.5 ± 3.3 | -194.2 ± 9.9 | -12.25 ± 0.20 |
| GCACA**PP**TGA | CGTGT**ZZ**ACT | -76.5 ± 3.6 | -203.4 ± 10.7 | -13.44 ± 0.25 |
| GCACA**ZP**GGA | CGTGT**PZ**CCT | -80.3 ± 3.5 | -212.5 ± 10.3 | -14.41 ± 0.27 |
| GCACC**PZ**TGA | CGTGG**ZP**ACT | -74.6 ± 2.8 | -195.1 ± 8.3 | -14.10 ± 0.21 |
| GCACG**PZ**GAA | CGTGC**ZP**CTT | -75.8 ± 4.5 | -198.8 ± 13.3 | -14.12 ± 0.35 |
| GCACT**P**ATGA | CGTGA**Z**TACT | -70.3 ± 3.2 | -191.1 ± 9.9 | -11.03 ± 0.14 |
| GCACT**P**TTGA | CGTGA**Z**AACT | -66.2 ± 2.6 | -178.0 ± 7.9 | -11.04 ± 0.13 |
| GCACT**ZP**GAA | CGTGA**PZ**CTT | -78.4 ± 4.5 | -208.7 ± 13.6 | -13.67 ± 0.33 |
| GCACT**ZZ**GAA | CGTGA**PP**CTT | -67.9 ± 3.7 | -178.2 ± 11.2 | -12.67 ± 0.25 |
| **P**CACAGATGA | **Z**GTGTCTACT | -70.0 ± 2.3 | -190.4 ± 7.0 | -10.94 ± 0.10 |
| **Z**CACAGATGA | **P**GTGTCTACT | -66.2 ± 2.3 | -180.8 ± 7.3 | -10.18 ± 0.08 |

Table S2. The nearest neighbor stack parameters for P-Z pairs with and without $(ZGCATGCP)_2$, which we identified as an outlier. These fits include the terminal P-Z penalty, which we determined to not be required.

| Parameter: | Including $(ZGCATGCP)_2$<br>$\Delta G°_{37}$ (kcal/mol) | Excluding $(ZGCATGCP)_2$<br>$\Delta G°_{37}$ (kcal/mol) |
|---|---|---|
| TP<br>AZ | -1.29 ± 0.17 | -1.45 ± 0.15 |
| AP<br>TZ | -1.60 ± 0.19 | -1.85 ± 0.16 |
| AZ<br>TP | -1.81 ± 0.21 | -1.45 ± 0.19 |
| TZ<br>AP | -1.83 ± 0.18 | -1.50 ± 0.17 |
| GZ<br>CP | -2.06 ± 0.20 | -1.75 ± 0.18 |
| GP<br>CZ | -2.03 ± 0.19 | -2.27 ± 0.16 |
| PZ<br>ZP | -2.12 ± 0.37 | -1.58 ± 0.33 |
| CZ<br>GP | -2.33 ± 0.16 | -2.08 ± 0.14 |
| PP<br>ZZ | -2.45 ± 0.17 | -2.36 ± 0.14 |
| CP<br>GZ | -2.35 ± 0.17 | -2.76 ± 0.17 |
| ZP<br>PZ | -2.70 ± 0.39 | -3.33 ± 0.35 |
| P end penalty<br>Z | 0.33 ± 0.21 | 0.03± 0.19 |

Table S3. The full set of duplexes used to fit P-Z pairs and the residuals of the fits.

| Sequence 1 5′ -> 3′ | Sequence 2 3′ -> 5′ | Source† | Experimental $\Delta G°_{37}$ (kcal/mol) | Experimental $\Delta G°_{37}$ for P-Z stacks (kcal/mol) | Fit $\Delta G°_{37}$ for P-Z stacks (kcal/mol) | Residual as (Experimental $\Delta G°_{37}$) – (Fit $\Delta G°_{37}$) (kcal/mol) |
|---|---|---|---|---|---|---|
| $CPGATCZG_2$ | | A | -10.52 | -9.45 | -9.68 | 0.23 |
| $CPPATZZG_2$ | | A | -12.07 | -13.60 | -13.20 | -0.40 |
| $CZACGTPG_2$ | | A | -10.23 | -7.66 | -7.05 | -0.61 |
| $CZCATGPG_2$ | | A | -9.87 | -8.40 | -8.71 | 0.31 |
| $CZTCGAPG_2$ | | A | -10.4 | -8.03 | -7.85 | -0.18 |
| $GACPZGTC_2$ | | A | -10.38 | -7.41 | -7.10 | -0.31 |
| $GACZPGTC_2$ | | A | -10.79 | -7.82 | -7.50 | -0.32 |
| $GAPATZTC_2$ | | A | -7.96 | -6.89 | -6.67 | -0.22 |
| $GAZATPTC_2$ | | A | -7.13 | -6.06 | -5.77 | -0.29 |
| $GAZCGPTC_2$ | | A | -9.92 | -7.55 | -7.44 | -0.11 |
| $GAZTAPTC_2$ | | A | -7.23 | -6.46 | -6.57 | 0.11 |
| $GAZZPPTC_2$ | | A | -11.87 | -11.70 | -10.93 | -0.77 |
| $GCAPZTGC_2$ | | A | -10.48 | -5.51 | -5.27 | -0.24 |
| $GCAZPTGC_2$ | | A | -10.99 | -6.02 | -6.23 | 0.21 |
| $GCPATZGC_2$ | | A | -11.46 | -8.59 | -8.50 | -0.09 |
| $GCPTAZGC_2$ | | A | -10.57 | -8.00 | -8.40 | 0.40 |
| $GCTPZAGC_2$ | | A | -8.83 | -4.26 | -4.47 | 0.21 |
| $GCTZPAGC_2$ | | A | -9.73 | -5.16 | -6.32 | 1.16 |
| $GGAPZTCC_2$ | | B | -9.65 | -5.88 | -5.27 | -0.61 |
| $GGAPZTCC_2$ | | A | -8.63 | -4.86 | -5.27 | 0.41 |
| $GGAZPTCC_2$ | | B | -10.1 | -6.33 | -6.23 | -0.10 |
| $GGAZPTCC_2$ | | A | -9.66 | -5.89 | -6.23 | 0.34 |
| $GGZATPCC_2$ | | A | -7.93 | -5.86 | -6.37 | 0.51 |
| $GGZTAPCC_2$ | | A | -8.38 | -6.61 | -7.17 | 0.56 |
| $GPACGTZC_2$ | | B | -10.5 | -7.93 | -7.54 | -0.39 |
| $GPACGTZC_2$ | | A | -9.51 | -6.94 | -7.54 | 0.60 |
| $GPCATGZC_2$ | | A | -8.59 | -7.12 | -8.04 | 0.92 |
| $GTGPZCAC_2$ | | A | -10.28 | -6.91 | -6.14 | -0.77 |
| $GTGZPCAC_2$ | | A | -10.81 | -7.44 | -6.83 | -0.61 |
| $GTPPZZAC_2$ | | A | -8.13 | -7.76 | -9.17 | 1.41 |
| $GTZATPAC_2$ | | A | -7.78 | -6.51 | -5.87 | -0.64 |
| $GTZCGPAC_2$ | | A | -10.65 | -8.08 | -7.54 | -0.54 |
| $GZGATCPC_2$ | | A | -10.39 | -9.32 | -9.00 | -0.32 |
| $GZZATPPC_2$ | | A | -9.86 | -11.39 | -11.07 | -0.32 |
| $GZZTAPPC_2$ | | A | -9.9 | -11.73 | -11.87 | 0.14 |
| $PACTAGTZ_2$ | | A | -6.4 | -2.83 | -2.97 | 0.14 |
| $PGACGTCZ_2$ | | B | -10.2 | -5.03 | -4.15 | -0.88 |

| | | | | | | |
|---|---|---|---|---|---|---|
| $PGACGTCZ_2$ | | A | -9.05 | -3.88 | -4.15 | 0.27 |
| $PGCATGCZ_2$ | | A | -8.61 | -2.74 | -4.15 | 1.41 |
| $ZACTAGTP_2$ | | A | -6.33 | -2.76 | -2.90 | 0.14 |
| GCACAGATCP | CGTGTCTAGZ | C | -11.71 | -2.31 | -2.77 | 0.45 |
| GCACAGTTTZ | CGTGTCAAAP | C | -10.61 | -1.31 | -1.49 | 0.17 |
| GCACAPCTGA | CGTGTZGACT | C | -12.25 | -3.55 | -3.59 | 0.03 |
| GCACAPPTGA | CGTGTZZACT | C | -13.44 | -6.04 | -5.64 | -0.40 |
| GCACAZPGGA | CGTGTPZCCT | C | -14.4 | -6.70 | -6.86 | 0.16 |
| GCACCPZTGA | CGTGGZPACT | C | -14.1 | -6.40 | -6.19 | -0.21 |
| GCACGPZGAA | CGTGCZPCTT | C | -14.12 | -6.52 | -6.62 | 0.10 |
| GCACTPATGA | CGTGAZTACT | C | -11.02 | -2.92 | -2.93 | 0.01 |
| GCACTPTTGA | CGTGAZAACT | C | -11.04 | -2.84 | -2.89 | 0.05 |
| GCACTZPGAA | CGTGAPZCTT | C | -13.67 | -6.97 | -6.91 | -0.05 |
| GCACTZZGAA | CGTGAPPCTT | C | -12.67 | -5.97 | -6.60 | 0.63 |
| GCCAPTTAA | CGGTZAATT | B | -9.14 | -3.04 | -3.29 | 0.25 |
| GCCAPTTAA | CGGTZAATT | A | -9.01 | -2.91 | -3.29 | 0.38 |
| GCZAGTTAA | CGPTCAATT | B | -9.38 | -3.88 | -3.52 | -0.36 |
| GCZAGTTAA | CGPTCAATT | A | -9.19 | -3.69 | -3.52 | -0.17 |
| GZCAGTTAA | CPGTCAATT | A | -9.35 | -4.55 | -4.02 | -0.53 |
| GZCAGTTAA | CPGTCAATT | B | -9.23 | -4.43 | -4.02 | -0.41 |
| GZZAGTTAA | CPPTCAATT | B | -9.59 | -6.29 | -5.54 | -0.76 |
| GZZAGTTAA | CPPTCAATT | A | -8.45 | -5.15 | -5.54 | 0.39 |
| PCACAGATGA | ZGTGTCTACT | C | -10.94 | -2.24 | -1.74 | -0.50 |
| ZCACAGATGA | PGTGTCTACT | C | -10.18 | -1.48 | -2.28 | 0.80 |

[†]Sources of data are A, Hoshika et al.[15]; B, Wang et al.[16]; C, this work.

Table S4. Optical melting data for duplexes designed to determine G-Z stack nearest neighbor parameters. Eleven are new sequences for this work. Uncertainties are the fit uncertainties from simultaneous fitting to melting experiments done at two concentrations as described in the Materials and Methods.

| Sequence 1 5′->3′ | Sequence 2 3′->5′ | ΔH° (kcal/mol) | ΔS° (cal mol$^{-1}$ K$^{-1}$) | ΔG°$_{37}$ (kcal/mol) | Source†/ comment |
|---|---|---|---|---|---|
| AGCCAGTTAZ | TCGGTCAATG | -63.6 ± 1.2 | -171.9 ± 3.7 | -10.27 ± 0.09 | This work |
| GCCACGZAAG | CGGTGZGTTC | -48.5 ± 2.0 | -128.3 ± 6.3 | -8.69 ± 0.06 | This work |
| GCCAGAZAAG | CGGTZTGTTC | -52.1 ± 1.9 | -140.0 ± 5.8 | -8.68 ± 0.06 | This work; duplicate runs |
| GCCAGAZAAG | CGGTZTGTTC | -50.7 ± 1.8 | -135.9 ± 5.5 | -8.54 ± 0.05 | |
| GCCAGCGPAG | CGGTCGZZTC | -63.9 ± 7.2 | -167.1 ± 21.8 | -12.09 ± 0.44 | This work |
| GCCAGTTAA | CGGTZAATT | -47.9 ± 2.4 | -129.2 ± 7.8 | -7.83 ± 0.07 | Wang, 5ZG |
| GCCAGZZPAG | CGGTCGGZTC | -63.6 ± 8.7 | -168.8 ± 26.6 | -11.25 ± 0.43 | This work |
| GCCAZZGAAG | CGGTGGZTTC | -41.9 ± 2.1 | -109.1 ± 6.5 | -8.05 ± 0.06 | This work |
| GCCGZTTAAG | CGGZPAATTC | -72.5 ± 1.8 | -197.2 ± 5.6 | -11.36 ± 0.10 | This work |
| GCZAGTTAA | CGGTCAATT | -49.6 ± 3.4 | -137.0 ± 10.9 | -7.07 ± 0.02 | Wang, 3ZG |
| GCZCGTTAAG | CGGGZAATTC | -50.0 ± 1.5 | -135.4 ± 4.8 | -7.98 ± 0.06 | This work |
| GGCGGTTAAG | CZGZCAATTC | -59.1 ± 4.5 | -164.5 ± 14.4 | -8.08 ± 0.02 | This work |
| GZCAGTTAA | CGGTCAATT | -52.8 ± 3.6 | -146.0 ± 11.9 | -7.52 ± 0.14 | Wang, 2ZG |
| GZPAGTTAAG | CGZTCAATTC | -63.8 ± 1.4 | -173.3 ± 4.4 | -10.01 ± 0.07 | This work |
| GZZAGTTAA | CGGTCAATT | -46.3 ± 2.8 | -128.4 ± 9.3 | -6.48 ± 0.07 | Wang, 2,3ZG |
| GZZZGTTAAG | CGGPCAATTC | -63.4 ± 3.5 | -171.0 ± 10.7 | -10.35 ± 0.15 | This work |

†Wang et al., 2017;[16] uncertainty estimates for ΔG°$_{37}$ from Wang, 2016.[55]

Table S5. The set of duplexes used to fit G-Z pair stacking nearest neighbor parameters and the residuals of the fits. Note that the residuals are 0 kcal/mol in most cases because there was no redundancy in the set of dinucleotides with G-Z pair stacks.

| Sequence 1 5′ -> 3′ | Sequence 2 3′ -> 5′ | Experimental $\Delta G°_{37}$ (kcal/mol) | Experimental $\Delta G°_{37}$ for G-Z stacks (kcal/mol) | Fit $\Delta G°_{37}$ for G-Z stacks (kcal/mol) | Residual as (Experimental $\Delta G°_{37}$) – (Fit $\Delta G°_{37}$) (kcal/mol) |
|---|---|---|---|---|---|
| AGCCAGTTAZ | TCGGTCAATG | -10.27 | -1.17 | -1.17 | 0.00 |
| GCCACGZAAG | CGGTGZGTTC | -8.69 | -1.49 | -1.49 | 0.00 |
| GCCAGAZAAG | CGGTZTGTTC | -8.68 | -2.88 | -2.81 | -0.07 |
| GCCAGAZAAG | CGGTZTGTTC | -8.54 | -2.74 | -2.81 | 0.07 |
| GCCAGCGPAG | CGGTCGZZTC | -12.09 | -2.30 | -2.30 | 0.00 |
| GCCAGTTAA | CGGTZAATT | -7.84 | -1.74 | -1.74 | 0.00 |
| GCCAGZZPAG | CGGTCGGZTC | -11.25 | -3.66 | -3.66 | 0.00 |
| GCCAZZGAAG | CGGTGGZTTC | -8.05 | -2.25 | -2.25 | 0.00 |
| GCCGZTTAAG | CGGZPAATTC | -11.36 | -3.61 | -3.61 | 0.00 |
| GCZAGTTAA | CGGTCAATT | -7.07 | -1.57 | -1.57 | 0.00 |
| GCZCGTTAAG | CGGGZAATTC | -7.98 | -3.88 | -3.88 | 0.00 |
| GGCGGTTAAG | CZGZCAATTC | -8.08 | -4.78 | -4.78 | 0.00 |
| GZCAGTTAA | CGGTCAATT | -7.61 | -2.81 | -2.81 | 0.00 |
| GZPAGTTAAG | CGZTCAATTC | -10.01 | -3.92 | -3.92 | 0.00 |
| GZZAGTTAA | CGGTCAATT | -6.50 | -3.20 | -3.20 | 0.00 |
| GZZZGTTAAG | CGGPCAATTC | -10.35 | -4.28 | -4.28 | 0.00 |

Table S6. Stacking enthalpy and entropy change parameters for stacks with P-Z pairs.

| Parameter | ΔH° (kcal/mol) | ΔS° (e.u.) |
|---|---|---|
| TP<br>AZ | -6.33 ± 1.45 | -14.83 ± 4.34 |
| AP<br>TZ | -8.22 ± 1.60 | -19.74 ± 4.77 |
| AZ<br>TP | -4.96 ± 1.63 | -11.04 ± 4.88 |
| TZ<br>AP | -7.36 ± 1.45 | -19.13 ± 4.34 |
| GZ<br>CP | -2.23 ± 1.61 | -0.98 ± 4.80 |
| PZ<br>ZP | -3.63 ± 3.23 | -6.67 ± 9.64 |
| GP<br>CZ | -9.23 ± 1.57 | -21.51 ± 4.69 |
| CZ<br>GP | -6.80 ± 1.27 | -14.92 ± 3.79 |
| PP<br>ZZ | -10.96 ± 1.32 | -27.45 ± 3.94 |
| CP<br>GZ | -12.32 ± 1.61 | -30.24 ± 4.80 |
| ZP<br>PZ | -15.43 ± 3.18 | -37.92 ± 9.51 |

Table S7A. The full set of duplexes used to fit P-Z stacking enthalpy change parameters and the residuals of the fits.

| Sequence 1 5′ -> 3′ | Sequence 2 3′ -> 5′ | Source† | Experimental $\Delta H°_{37}$ (kcal/mol) | Experimental $\Delta H°_{37}$ for P-Z stacks (kcal/mol) | Fit $\Delta H°_{37}$ for P-Z stacks (kcal/mol) | Residual as (Experimental $\Delta H°_{37}$) – (Fit $\Delta H°_{37}$) (kcal/mol) |
|---|---|---|---|---|---|---|
| $CPGATCZG_2$ | | A | -62.9 | -39.5 | -38.2 | -1.2 |
| $CPPATZZG_2$ | | A | -71.2 | -64.2 | -61.3 | -2.9 |
| $CZACGTPG_2$ | | A | -57.8 | -30.6 | -26.3 | -4.3 |
| $CZCATGPG_2$ | | A | -51.1 | -27.0 | -32.1 | 5.0 |
| $CZTCGAPG_2$ | | A | -61.1 | -34.3 | -30.0 | -4.2 |
| $GACPZGTC_2$ | | A | -57.7 | -24.7 | -28.3 | 3.6 |
| $GACZPGTC_2$ | | A | -57.0 | -24.0 | -29.0 | 5.1 |
| $GAPATZTC_2$ | | A | -59.9 | -36.5 | -31.1 | -5.3 |
| $GAZATPTC_2$ | | A | -45.7 | -22.3 | -22.6 | 0.3 |
| $GAZCGPTC_2$ | | A | -52.4 | -25.6 | -28.4 | 2.8 |
| $GAZTAPTC_2$ | | A | -46.0 | -22.6 | -26.4 | 3.8 |
| $GAZZPPTC_2$ | | A | -66.0 | -49.8 | -47.3 | -2.5 |
| GCACAGATCP | CGTGTCTAGZ | C | -75.4 | -9.0 | -12.3 | 3.3 |
| GCACAGTTTZ | CGTGTCAAAP | C | -72.4 | -6.0 | -7.4 | 1.3 |
| GCACAPCTGA | CGTGTZGACT | C | -72.5 | -15.2 | -10.4 | -4.7 |
| GCACAPPTGA | CGTGTZZACT | C | -76.5 | -27.0 | -24.1 | -2.9 |
| GCACAZPGGA | CGTGTPZCCT | C | -80.3 | -31.3 | -27.2 | -4.1 |
| GCACCPZTGA | CGTGGZPACT | C | -74.6 | -25.6 | -24.2 | -1.4 |
| GCACGPZGAA | CGTGCZPCTT | C | -75.8 | -25.1 | -25.2 | 0.1 |
| GCACTPATGA | CGTGAZTACT | C | -70.3 | -14.3 | -13.7 | -0.6 |
| GCACTPTTGA | CGTGAZAACT | C | -66.2 | -9.8 | -11.3 | 1.4 |
| GCACTZPGAA | CGTGAPZCTT | C | -78.4 | -30.5 | -29.6 | -0.9 |
| GCACTZZGAA | CGTGAPPCTT | C | -67.9 | -20.0 | -30.6 | 10.6 |
| $GCAPZTGC_2$ | | A | -60.2 | -23.7 | -20.1 | -3.7 |
| $GCAZPTGC_2$ | | A | -64.6 | -28.2 | -25.4 | -2.8 |
| GCCAPTTAA | CGGTZAATT | B | -56.2 | -9.9 | -13.2 | 3.3 |
| GCCAPTTAA | CGGTZAATT | A | -58.5 | -12.2 | -13.2 | 1.0 |
| $GCPATZGC_2$ | | A | -67.1 | -40.5 | -39.4 | -1.1 |
| $GCPTAZGC_2$ | | A | -61.9 | -35.3 | -34.6 | -0.7 |
| $GCTPZAGC_2$ | | A | -46.0 | -11.0 | -16.3 | 5.3 |
| $GCTZPAGC_2$ | | A | -55.7 | -20.7 | -30.1 | 9.5 |
| GCZAGTTAA | CGPTCAATT | B | -58.5 | -12.5 | -13.1 | 0.7 |
| GCZAGTTAA | CGPTCAATT | A | -65.1 | -19.1 | -13.1 | -6.0 |

| | | | | | | |
|---|---|---|---|---|---|---|
| GGAPZTCC$_2$ | | A | -45.3 | -13.1 | -20.1 | 7.0 |
| GGAPZTCC$_2$ | | B | -58.1 | -25.9 | -20.1 | -5.8 |
| GGAZPTCC$_2$ | | A | -55.0 | -22.8 | -25.4 | 2.6 |
| GGAZPTCC$_2$ | | B | -60.4 | -28.2 | -25.4 | -2.8 |
| GGZATPCC$_2$ | | A | -36.5 | -13.5 | -17.1 | 3.7 |
| GGZTAPCC$_2$ | | A | -36.4 | -13.4 | -20.9 | 7.5 |
| GPACGTZC$_2$ | | A | -57.3 | -30.1 | -33.2 | 3.1 |
| GPACGTZC$_2$ | | B | -67.3 | -40.1 | -33.2 | -6.9 |
| GPCATGZC$_2$ | | A | -39.7 | -15.7 | -22.9 | 7.3 |
| GTGPZCAC$_2$ | | A | -59.2 | -25.6 | -22.1 | -3.5 |
| GTGZPCAC$_2$ | | A | -57.5 | -23.9 | -19.9 | -4.0 |
| GTPPZZAC$_2$ | | A | -56.4 | -39.8 | -38.2 | -1.5 |
| GTZATPAC$_2$ | | A | -52.7 | -28.9 | -27.4 | -1.5 |
| GTZCGPAC$_2$ | | A | -62.3 | -35.1 | -33.2 | -1.9 |
| GZCAGTTAA | CPGTCAATT | B | -58.5 | -13.8 | -11.5 | -2.3 |
| GZCAGTTAA | CPGTCAATT | A | -71.6 | -26.9 | -11.5 | -15.4 |
| GZGATCPC$_2$ | | A | -56.4 | -33.0 | -29.1 | -3.9 |
| GZZAGTTAA | CPPTCAATT | B | -58.3 | -22.1 | -19.5 | -2.5 |
| GZZAGTTAA | CPPTCAATT | A | -58.7 | -22.5 | -19.5 | -2.9 |
| GZZATPPC$_2$ | | A | -43.4 | -36.4 | -39.0 | 2.6 |
| GZZTAPPC$_2$ | | A | -46.7 | -39.7 | -42.8 | 3.2 |
| PACTAGTZ$_2$ | | A | -52.2 | -12.8 | -14.7 | 1.9 |
| PCACAGATGA | ZGTGTCTACT | C | -70.0 | -7.1 | -2.2 | -4.9 |
| PGACGTCZ$_2$ | | A | -54.7 | -11.1 | -13.6 | 2.5 |
| PGACGTCZ$_2$ | | B | -69.3 | -25.7 | -13.6 | -12.1 |
| PGCATGCZ$_2$ | | A | -42.8 | 0.8 | -13.6 | 14.4 |
| ZACTAGTP$_2$ | | A | -51.7 | -12.3 | -12.7 | 0.4 |
| ZCACAGATGA | PGTGTCTACT | C | -66.2 | -3.3 | -9.2 | 5.9 |

†Sources of data are A, Hoshika et al.[15]; B, Wang et al.[16]; C, this work.

Table S7B. The full set of duplexes used to fit P-Z stacking entropy change parameters and the residuals of the fits.

| Sequence 1 5′ -> 3′ | Sequence 2 3′ -> 5′ | Source† | Experimental $\Delta S°_{37}$ (e.u.) | Experimental $\Delta S°_{37}$ for P-Z stacks (e.u.) | Fit $\Delta S°_{37}$ for P-Z stacks (e.u.) | Residual as (Experimental $\Delta S°_{37}$) – (Fit $\Delta S°_{37}$) (e.u) |
|---|---|---|---|---|---|---|
| $CPGATCZG_2$ | | A | -168.72 | -96.82 | -90.31 | -6.52 |
| $CPPATZZG_2$ | | A | -190.65 | -163.15 | -153.64 | -9.51 |
| $CZACGTPG_2$ | | A | -153.22 | -74.12 | -59.49 | -14.62 |
| $CZCATGPG_2$ | | A | -132.77 | -59.87 | -72.86 | 12.98 |
| $CZTCGAPG_2$ | | A | -163.31 | -84.61 | -69.33 | -15.28 |
| $GACPZGTC_2$ | | A | -152.57 | -56.27 | -67.14 | 10.87 |
| $GACZPGTC_2$ | | A | -148.83 | -52.53 | -67.75 | 15.22 |
| $GAPATZTC_2$ | | A | -167.31 | -95.41 | -77.76 | -17.65 |
| $GAZATPTC_2$ | | A | -124.20 | -52.30 | -51.74 | -0.56 |
| $GAZCGPTC_2$ | | A | -136.97 | -58.27 | -65.10 | 6.84 |
| $GAZTAPTC_2$ | | A | -125.00 | -52.20 | -61.57 | 9.37 |
| $GAZZPPTC_2$ | | A | -174.37 | -122.87 | -114.90 | -7.97 |
| GCACAGATCP | CGTGTCTAGZ | C | -205.30 | -21.60 | -30.24 | 8.64 |
| GCACAGTTTZ | CGTGTCAAAP | C | -199.38 | -15.48 | -19.13 | 3.65 |
| GCACAPCTGA | CGTGTZGACT | C | -194.19 | -28.99 | -20.73 | -8.26 |
| GCACAPPTGA | CGTGTZZACT | C | -203.39 | -59.19 | -58.24 | -0.95 |
| GCACAZPGGA | CGTGTPZCCT | C | -212.54 | -71.14 | -63.88 | -7.26 |
| GCACCPZTGA | CGTGGZPACT | C | -195.12 | -53.72 | -56.65 | 2.93 |
| GCACGPZGAA | CGTGCZPCTT | C | -198.75 | -51.45 | -58.41 | 6.96 |
| GCACTPATGA | CGTGAZTACT | C | -191.06 | -28.16 | -33.96 | 5.80 |
| GCACTPTTGA | CGTGAZAACT | C | -178.01 | -14.21 | -25.87 | 11.66 |
| GCACTZPGAA | CGTGAPZCTT | C | -208.72 | -67.62 | -71.97 | 4.34 |
| GCACTZZGAA | CGTGAPPCTT | C | -178.21 | -37.11 | -76.82 | 39.71 |
| $GCAPZTGC_2$ | | A | -160.15 | -58.85 | -46.16 | -12.69 |
| $GCAZPTGC_2$ | | A | -172.69 | -71.39 | -60.00 | -11.39 |
| GCCAPTTAA | CGGTZAATT | B | -151.66 | -13.66 | -30.79 | 17.13 |
| GCCAPTTAA | CGGTZAATT | A | -159.57 | -21.57 | -30.79 | 9.22 |
| $GCPATZGC_2$ | | A | -179.40 | -103.10 | -98.74 | -4.36 |
| $GCPTAZGC_2$ | | A | -165.34 | -88.14 | -82.55 | -5.59 |
| $GCTPZAGC_2$ | | A | -119.85 | -21.95 | -36.33 | 14.38 |
| $GCTZPAGC_2$ | | A | -148.06 | -50.16 | -76.18 | 26.02 |
| GCZAGTTAA | CGPTCAATT | B | -158.24 | -19.44 | -29.75 | 10.30 |
| GCZAGTTAA | CGPTCAATT | A | -180.27 | -41.47 | -29.75 | -11.72 |

| | | | | | | |
|---|---|---|---|---|---|---|
| $GGAPZTCC_2$ | | A | -118.23 | -26.93 | -46.16 | 19.23 |
| $GGAPZTCC_2$ | | B | -156.21 | -64.91 | -46.16 | -18.76 |
| $GGAZPTCC_2$ | | A | -146.03 | -54.73 | -60.00 | 5.27 |
| $GGAZPTCC_2$ | | B | -162.18 | -70.88 | -60.00 | -10.88 |
| $GGZATPCC_2$ | | A | -91.96 | -24.66 | -31.63 | 6.97 |
| $GGZTAPCC_2$ | | A | -90.18 | -21.98 | -41.46 | 19.48 |
| $GPACGTZC_2$ | | A | -154.09 | -74.99 | -81.29 | 6.30 |
| $GPACGTZC_2$ | | B | -183.14 | -104.04 | -81.29 | -22.75 |
| $GPCATGZC_2$ | | A | -100.15 | -27.25 | -44.99 | 17.74 |
| $GTGPZCAC_2$ | | A | -157.57 | -60.27 | -49.69 | -10.58 |
| $GTGZPCAC_2$ | | A | -150.54 | -53.24 | -39.88 | -13.36 |
| $GTPPZZAC_2$ | | A | -155.47 | -103.57 | -91.23 | -12.34 |
| $GTZATPAC_2$ | | A | -144.83 | -72.53 | -67.92 | -4.61 |
| $GTZCGPAC_2$ | | A | -166.37 | -87.27 | -81.29 | -5.98 |
| GZCAGTTAA | CPGTCAATT | B | -158.86 | -21.76 | -22.49 | 0.73 |
| GZCAGTTAA | CPGTCAATT | A | -200.55 | -63.45 | -22.49 | -40.95 |
| $GZGATCPC_2$ | | A | -148.35 | -76.45 | -62.44 | -14.01 |
| GZZAGTTAA | CPPTCAATT | B | -156.89 | -42.49 | -43.26 | 0.77 |
| GZZAGTTAA | CPPTCAATT | A | -161.86 | -47.46 | -43.26 | -4.19 |
| $GZZATPPC_2$ | | A | -108.14 | -80.64 | -86.53 | 5.89 |
| $GZZTAPPC_2$ | | A | -118.49 | -90.09 | -96.36 | 6.27 |
| $PACTAGTZ_2$ | | A | -147.67 | -32.47 | -38.26 | 5.79 |
| PCACAGATGA | ZGTGTCTACT | C | -190.43 | -7.03 | -0.98 | -6.05 |
| $PGACGTCZ_2$ | | A | -147.19 | -23.69 | -29.83 | 6.15 |
| $PGACGTCZ_2$ | | B | -190.55 | -67.05 | -29.83 | -37.22 |
| $PGCATGCZ_2$ | | A | -110.08 | 11.62 | -29.83 | 41.46 |
| $ZACTAGTP_2$ | | A | -146.28 | -31.08 | -29.66 | -1.42 |
| ZCACAGATGA | PGTGTCTACT | C | -180.75 | 2.65 | -21.51 | 24.16 |

†Sources of data are A, Hoshika et al.[15]; B, Wang et al.[16]; C, this work.

Table S8. Optical melting data for eight systems with dangling ends (this work).

| Sequence 1 5′->3′ | Sequence 2 3′->5′ | ΔH° (kcal/mol) | ΔS° (cal $mol^{-1}$ $K^{-1}$) | $\Delta G°_{37}$ (kcal/mol) |
|---|---|---|---|---|
| **P**CACAGATGA | GTGTCTACT | -65.4 ± 2.4 | -181.5 ± 7.6 | -9.07 ± 0.05 |
| CACAGATGA | **P**GTGTCTACT | -61.4 ± 3.5 | -170.2 ± 11.0 | -8.63 ± 0.05 |
| **Z**CACAGATGA | GTGTCTACT | -64.3 ± 3.3 | -177.2 ± 10.3 | -9.34 ± 0.08 |
| CACAGATGA | **Z**GTGTCTACT | -64.3 ± 3.3 | -177.2 ± 10.3 | -9.34 ± 0.08 |
| GCACAGTTT**Z** | CGTGTCAAA | -60.4 ± 2.8 | -164.2 ± 8.8 | -9.47 ± 0.08 |
| GCACAGTTT | CGTGTCAAA**Z** | -72.5 ± 2.5 | -201.1 ± 7.7 | -10.10 ± 0.08 |
| GCACAGTTT**P** | CGTGTCAAA | -61.2 ± 1.2 | -167.5 ± 3.7 | -9.21 ± 0.03 |
| GCACAGTTT | CGTGTCAAA**P** | -70.8 ± 1.7 | -196.3 ± 5.3 | -9.93 ± 0.05 |

Table S9. Optical melting data for thirteen systems with terminal mismatches (this work).

| Sequence 1 5′->3′ | Sequence 2 3′->5′ | ΔH° (kcal/mol) | ΔS° (cal mol$^{-1}$ K$^{-1}$) | $\Delta G^\circ_{37}$ (kcal/mol) |
|---|---|---|---|---|
| **P**CACAGATGA | **P**GTGTCTACT | -61.3 ± 1.6 | -168.2 ± 4.9 | -9.14 ± 0.03 |
| **Z**CACAGATGA | **Z**GTGTCTACT | -64.1 ± 2.4 | -176.2 ± 7.4 | -9.46 ± 0.06 |
| GCACAGTTT**Z** | CGTGTCAAA**Z** | -64.5 ± 2.3 | -175.2 ± 7.3 | -10.21 ± 0.08 |
| GCACAGTTT**P** | CGTGTCAAA**P** | -62.6 ± 2.7 | -170.3 ± 8.4 | -9.81 ± 0.08 |
| G**Z**CAGTTGAA | G**P**GTCAACTT | -61.9 ± 2.9 | -167.9 ± 9.0 | -9.81 ± 0.10 |
| G**Z**CAGTTGAA | **P**GTCAACTT | -61.8 ± 3.2 | -168.8 ± 9.9 | -9.45 ± 0.08 |
| A**Z**CAGTTGAA | A**P**GTCAACTT | -65.0 ± 1.7 | -178.6 ± 5.4 | -9.56 ± 0.05 |
| A**Z**CAGTTGAA | **P**GTCAACTT | -65.0 ± 1.7 | -178.4 ± 5.5 | -9.62 ± 0.05 |
| C**Z**CAGTTGAA | C**P**GTCAACTT | -62.7 ± 2.2 | -172.3 ± 6.9 | -9.28 ± 0.06 |
| C**Z**CAGTTGAA | **P**GTCAACTT | -61.8 ± 2.0 | -169.9 ± 6.4 | -9.11 ± 0.05 |
| G**Z**CAGTTGAA | A**P**GTCAACTT | -61.7 ± 1.5 | -168.3 ± 4.7 | -9.52 ± 0.04 |
| T**Z**CAGTTGAA | T**P**GTCAACTT | -61.6 ± 1.3 | -168.1 ± 4.1 | -9.43 ± 0.03 |
| T**Z**CAGTTGAA | **P**GTCAACTT | -63.5 ± 1.5 | -174.7 ± 4.8 | -9.36 ± 0.04 |

Table S10. Optical melting data for systems with single mismatches (from Wang et al., 2017, Table 2)[16] and two tandem mismatches (this work). NA: $T_m$ between 5 °C and 15 °C, so ΔH° and ΔS° are not reliable. Four 9-bp tandem mismatch duplexes from Wang et al. did not show melting transitions with $T_m$ > 5 °C, not shown here.

| Sequence 1 5′->3′ | Sequence 2 3′->5′ | ΔH° (kcal/mol) | ΔS° (cal mol$^{-1}$ K$^{-1}$) | ΔG°$_{37}$ (kcal/mol) | Source† |
|---|---|---|---|---|---|
| GCCA**P**TTAA | CGGT**C**AATT | -58.6 ± 2.6 | -171.0 ± 8.7 | -5.5 ± 0.1 | Wang |
| GC**C**AGTTAA | CG**P**TCAATT | -52.6 ± 2.4 | -151.4 ± 8.2 | -5.6 ± 0.1 | Wang |
| G**C**CAGTTAA | C**P**GTCAATT | -49.2 ± 2.8 | -139.8 ± 9.3 | -5.8 ± 0.1 | Wang |
| GCCA**P**TTAA | CGGT**T**AATT | -61.3 ± 3.8 | -183.2 ± 12.7 | -4.5 ± 0.2 | Wang |
| GC**T**AGTTAA | CG**P**TCAATT | NA | NA | -4.6 ± 0.1 | Wang |
| G**T**CAGTTAA | C**P**GTCAATT | -45.4 ± 3.4 | -128.1 ± 11.2 | -5.7 ± 0.1 | Wang |
| GCCA**A**TTAA | CGGT**Z**AATT | NA | NA | -4.7 ± 0.2 | Wang |
| GC**Z**AGTTAA | CG**A**TCAATT | NA | NA | NA | Wang |
| G**Z**CAGTTAA | C**A**GTCAATT | -41.3 ± 3.4 | -115.7 ± 11.3 | -5.4 ± 0.1 | Wang |
| GCACT**ZZ**GAA | CGTGA**ZZ**CTT | -27.3 ± 8.5 | -66.3 ± 27.4 | -6.72 ± 0.02 | This Work |
| GCACT**PP**GAA | CGTGA**PP**CTT | -24.4 ± 3.7 | -60.1 ± 13.6 | -5.72 ± 0.50 | This Work |

†Source of prior experiments is Wang et al.[16]

Table S11A. 5′ Dangling ends on a Z-P pair as compared to C-G and T-A pairs.

| 5' dangling end on Z-P (kcal/mol) | | 5' dangling end on C-G (kcal/mol) | | 5' dangling end on T-A (kcal/mol) | |
|---|---|---|---|---|---|
| 5′GZ<br>3′ P | -0.16 | 5′GC<br>3′ G | -0.7 | 5′GT<br>3′ A | -0.5 |
| 5′AZ<br>3′ P | -0.33 | 5′AC<br>3′ G | -0.9 | 5′AT<br>3′ A | -0.5 |
| 5′CZ<br>3′ P | 0.18 | 5′CC<br>3′ G | -0.5 | 5′CT<br>3′ A | -0.2 |
| 5′TZ<br>3′ P | -0.07 | 5′TC<br>3′ G | -0.6 | 5′TT<br>3′ A | -0.3 |

Table S11B. P and Z dangling ends compared to canonical nucleotides.

| Dangling end (kcal/mol) | | Comparable dangling end (kcal/mol) | | Comparable dangling end (kcal/mol) | |
|---|---|---|---|---|---|
| 5′PC<br>3′ G | -0.37 | 5′AC<br>3′ G | -0.9 | 5′GC<br>3′ G | -0.7 |
| 5′PA<br>3′ T | -0.63 | 5′AA<br>3′ T | -0.5 | 5′GA<br>3′ T | -0.6 |
| 5′ C<br>3′PG | 0.07 | 5′ C<br>3′AG | -0.4 | 5′ C<br>3′GG | -0.4 |
| 5′ A<br>3′PT | 0.09 | 5′ A<br>3′AT | -0.2 | 5′ A<br>3′GT | -0.2 |
| 5′ZC<br>3′ G | -0.64 | 5′CC<br>3′ G | -0.5 | 5′TC<br>3′ G | -0.6 |
| 5′ZA<br>3′ T | -0.80 | 5′CA<br>3′ T | -0.2 | 5′TA<br>3′ T | -0.3 |
| 5′ C<br>3′ZG | -0.11 | 5′ C<br>3′CG | -0.2 | 5′ C<br>3′TG | -0.8 |
| 5′ A<br>3′ZT | -0.17 | 5′ A<br>3′CT | -0.2 | 5′ A<br>3′TT | -0.2 |

Table S12. Stability of PP or ZZ terminal mismatches. The stabilities of analogous purine-purine or pyrimidine-pyrimidine mismatches are shown for comparison.

| Terminal Mismatch | Stability ($\Delta G°_{37}$; kcal/mol) | Analogous Mismatch | Stability ($\Delta G°_{37}$; kcal/mol) | Analogous Mismatch | Stability ($\Delta G°_{37}$; kcal/mol) |
|---|---|---|---|---|---|
| 5′GP<br>3′CP | -0.44 | 5′GG<br>3′CG | -1.0 | 5′GA<br>3′CA | -1.0 |
| 5′TP<br>3′AP | -0.51 | 5′TG<br>3′AG | -0.4 | 5′TA<br>3′AA | -0.6 |
| 5′GZ<br>3′CZ | -0.76 | 5′GC<br>3′CC | -0.6 | 5′GT<br>3′CT | -0.9 |
| 5′TZ<br>3′AZ | -0.91 | 5′TC<br>3′AC | -0.2 | 5′TT<br>3′AT | -0.3 |

Table S13. Stability of mismatches on terminal Z-P pairs. The analogous stabilities for mismatches on C-G terminal pairs are shown for comparison.

| Terminal Mismatch | Stability ($\Delta G^\circ_{37}$; kcal/mol) | Analogous Mismatch | Stability ($\Delta G^\circ_{37}$; kcal/mol) |
|---|---|---|---|
| 5′GZ<br>3′GP | -0.52 | 5′GC<br>3′GG | -1.0 |
| 5′AZ<br>3′AP | -0.27 | 5′AC<br>3′AG | -1.0 |
| 5′CZ<br>3′CP | 0.01 | 5′CC<br>3′CG | -0.6 |
| 5′GZ<br>3′AP | -0.23 | 5′GC<br>3′AG | -1.0 |
| 5′TZ<br>3′TP | -0.14 | 5′TC<br>3′TG | -0.9 |

Table S14. Single mismatch (1×1 internal loop) folding free energy changes.

| Duplex | Loop Stability (kcal/mol) |
|---|---|
| 5'GCCAPTTAA3'<br>3'CGGTCAATT5' | 0.6 |
| 5'GCCAGTTAA3'<br>3'CGPTCAATT5' | -0.1 |
| 5'GCCAGTTAA3'<br>3'CPGTCAATT5' | -1.0 |
| 5'GCCAGTTAA3'<br>3'CGGTTAATT5' | 0.7 |
| 5'GCTAGTTAA3'<br>3'CGGTCAATT5' | 0.1 |
| 5'GTCAGTTAA3'<br>3'CGGTCAATT5' | -1.1 |
| 5'GCCAPTTAA3'<br>3'CGGTTAATT5' | 1.6 |
| 5'GCTAGTTAA3'<br>3'CGPTCAATT5' | 0.9 |
| 5'GTCAGTTAA3'<br>3'CPGTCAATT5' | -0.9 |
| 5'GZCAGTTAA3'<br>3'CAGTCAATT5' | -0.6 |
| 5'GCCAGTTAA3'<br>3'CAGTCAATT5' | -0.4 |

Table S15. Tandem mismatch (2×2 internal loop) folding free energy changes.

| Motif | Motif Stability (kcal/mol) | Analogous Motif 1 | Analogous Motif 1 Stability (kcal/mol) | Analogous Motif 2 | Analogous Motif 2 Stability (kcal/mol) |
|---|---|---|---|---|---|
| 5′TZZG3′<br>3′AZZC5′ | -0.42 | 5′TCCG3′<br>3′ACCC5′ | 2.3 | 5′TTTG3′<br>3′ATTC5′ | 1.6 |
| 5′TPPG3′<br>3′APPC5′ | 0.58 | 5′TGGG3′<br>3′AGGC5′ | 1.6 | 5′TAAG3′<br>3′AAAC5′ | 1.5 |

Table S16. DNAzyme designs with or without P-Z pairs.

| | Canonical DNA | | DNA Including P-Z | |
|---|---|---|---|---|
| Design # | Time (s) | NED | Time (s) | NED |
| 1 | 1573.3 | 0.064 | 1576.8 | 0.040 |
| 2 | 4634.0 | 0.074 | 1375.8 | 0.031 |
| 3 | 2858.7 | 0.071 | 1769.2 | 0.046 |
| 4 | 7342.7 | 0.041 | 1492.1 | 0.036 |
| 5 | 2313.9 | 0.104 | 1654.5 | 0.037 |
| 6 | 4067.5 | 0.138 | 1589.3 | 0.044 |
| 7 | 3378.9 | 0.078 | 1507.3 | 0.044 |
| 8 | 2049.3 | 0.100 | 1826.9 | 0.054 |
| 9 | 1461.8 | 0.203 | 1633.5 | 0.054 |
| 10 | 2636.4 | 0.174 | 1517.2 | 0.033 |
| Mean: | 3231.6 | 0.105 | 1594.2 | 0.042 |
| Standard Deviation: | 1770.2 | | 134.1 | |
| Best: | | 0.041 | | 0.031 |
